# AIedes: a neural-network framework for modelling the distribution and abundance of *Aedes albopictus* in Europe

**DOI:** 10.64898/2026.05.14.722605

**Authors:** Indaco Biazzo, Lia Orfei, Ioannis Kioutsioukis, Lea Schuh, Céline Gossner, Olivier Briet, Peter V. Markov

**Author notes:** Corresponding authors *Email addresses:* (Indaco Biazzo), (Lea Schuh).

## Abstract

Modelling the distribution and abundance of invasive species requires the integration of heterogeneous ecological surveillance data with environmental covariates and reproducible computational workflows. *Aedes albopictus* is an invasive mosquito species whose ongoing geographical expansion across Europe, driven by climate change, trade, human mobility and urbanisation, is increasing both ecological and public-health concerns. Accurate mapping of its distribution and abundance is therefore essential for monitoring, forecasting and targeted vector-control strategies.

Here we present ***AI****edes*, a neural-network framework that predicts both the presence and weekly egg abundance of *Ae. albopictus* across Europe using climate variables and harmonised surveillance data. The model reproduces the large-scale spatial distribution of the species and captures fine-grained spatiotemporal variation in egg-laying activity. To support reproducible evaluation, the framework is trained and validated on newly harmonised, largescale surveillance and climate datasets, which are publicly available together with the model implementation and code. These resources provide a consistent benchmark for comparing vector distribution and abundance models and offer an extensible workflow for ecological forecasting of invasive mosquito species in other regions where comparable data are avail-able.

## 1. Introduction

The modelling of invasive species distributions and population dynamics is a central problem in ecological informatics, requiring the integration of heterogeneous surveillance records, environmental covariates and reproducible computational workflows. *Aedes albopictus* (Skuse, 1895), commonly known as the Asian tiger mosquito, is an invasive species that has considerably expanded its global presence over the past decades. Recent studies indicate that *Ae. albopictus* is now found in 126 countries and territories, on all continents except Antarctica [1], demonstrating ecological plasticity and adaptability to varying climatic conditions. By 2080, most countries in the world are projected to report *Ae. albopictus* [2]. Changes in climate appear to be an important factor for this geographical expansion [3], increasing the risk of mosquito-borne viral infections across a range of latitudes, posing a public health threat and a potential economic burden. The species continues its rapid expansion across Europe and parts of the Mediterranean Basin, recently establishing in Cyprus and Slovakia, and expanding its distribution in Austria, Belgium, France, Germany, Greece, Hungary, Portugal, Serbia, Slovenia, Spain, Switzerland, Tunisia and Türkiye [4]. The expansion of *Ae. albopictus* has been driven by the increasing availability of areas with suitable climate conditions for its establishment and by several biological and anthropogenic factors, including growing urbanisation, the species’ ability to enter diapause during winter, which enhances survival across diverse climatic regions, and the remarkable desiccation tolerance of its eggs, which supports persistence under dry conditions, while intensified global trade and human mobility have further facilitated its long-distance dispersal and colonisation of new areas [5].

*Ae. albopictus* is competent in the transmission of a number of arboviruses, including those causing dengue, chikungunya virus disease, and Zika virus disease [6]. The establishment of *Ae. albopictus* populations is a precondition for the local vector-borne transmission of these diseases, as clearly demonstrated by recent autochthonous cases of chikungunya and dengue in Southern Europe [7, 8].

Vector abundance, among other factors, contributes to transmission risk. However, abundance data are costly to obtain and are available for relatively few locations. Predictions of ovitrap-derived egg-laying activity may therefore contribute to entomological surveillance planning. They do not directly estimate disease risk, and their use for intervention decisions would require local validation and integration with epidemiological, entomological, and operational information.

Traditional approaches to predicting mosquito distribution typically rely on two types of models. The first type split the mosquito population into compartments based on stages in their life cycle (egg, larva, pupa, adult), and have been used extensively to model vector-host dynamics and disease transmission patterns [9, 10, 11, 12, 13, 14, 15]. These models provide mechanistic insights into epidemiological processes, but often require knowledge of biological and ecological parameters that may be difficult to estimate or may vary significantly across space, time and other circumstances, and these uncertainties may affect the accuracy of the models’ results. The second type are statistical models, typically mixed-effect models, such as generalised linear mixed models (GLMM) or generalised additive models (GAM) [16], and fixed-effect models, such as generalised linear models (GLM) [17, 18] and geographically weighted regression (GWR) [19]. According to a recent review of risk-mapping approaches for *Aedes*-borne viruses [20], the latter class of models enjoys by far the widest use. Specifically tailored for ecological and environmental applications, species distribution models (SDMs) are increasingly common statistical approaches to investigate the spread of mosquitoes, despite sub-optimal practices regarding quality of the response variable, predictor variables and model building [21]. Maximum entropy (MaxEnt) models are used in over 70% of mosquito SDM studies [22], yet these models often employ a large number of parameters whose inference can be challenging. It is often difficult to estimate them directly from data or to assign objective values based on literature or expert knowledge and they rely on over-simplistic assumptions that may miss the complex, non-linear interactions between environmental factors and mosquito behaviour.

Machine learning techniques have recently been employed to improve the prediction of mosquito species distribution and abundance, facilitated by the increasing availability of data for training. Ensemble approaches combining Random Forest and XGBoost classifiers have shown promising results for habitat suitability prediction [23]. Other ensemble modeling strategies have addressed model selection uncertainties and variability, by combining multiple statistical models with diverse machine learning algorithms to enhance prediction accuracy and reliability while minimizing the influence of individual model constraints [24]. Stacked machine learning models have demonstrated improved performance in capturing spatio-temporal dynamics of *Ae. albopictus* populations compared to mechanistic and correlative approaches used in isolation or aggregated via simple methods like averages and weighted averages [25]. However, many of these approaches rely on predefined sets of input features and may struggle to adapt to spatio-temporal variations in mosquito populations, particularly when modeling the complex ecological interactions that drive vector abundance patterns.

The application of neural networks to vector ecology could represent a complementary approach to traditional mechanistic and correlative methods [26]. Although they also require a choice of input variables, neural networks can automatically learn complex, hierarchical representations from raw environmental features without the need for extensive manual feature engineering [27]. This capability is particularly valuable in mosquito ecology, where abundance responses to environmental variables often exhibit threshold effects, counterintuitive non-linear interactions, and complex temporal dependencies on temperature and other variables [28, 29].

Recent studies have applied neural network approaches to mosquito modeling. One approach uses a neural network to generate forecasts of *Aedes aegypti* abundance from weather data, supporting vector surveillance [30]. In this case, the network is trained on synthetic data generated by a mechanistic simulation model, and once trained, it provides forecasts faster and at lower computational cost than running the simulation itself. Another study integrates neural networks within a mechanistic model, enabling data-driven estimation of key biological functions such as the relationship between temperature, precipitation and oviposition, within differential equations for population dynamics [31].

In this paper, we present ***AI****edes*, a data-driven neural-network framework for estimating the annual spatial distribution and weekly egg abundance of *Ae. albopictus* in Europe. Throughout this paper, abundance refers to ovitrap-derived egg abundance, that is the number of eggs collected per trap per week; it serves as a proxy for adult mosquito densities rather than a direct measure of them. The contribution of this work is threefold. First, we harmonise large-scale mosquito surveillance data with climate covariates at spatial and temporal resolutions suitable for ecological modelling. Second, we develop two complementary neuralnetwork models: ***AI****edes*-Classifier for broad-scale presence–absence prediction and ***AI****edes*Counter for weekly abundance prediction. Third, by using standard performance metrics, explicit validation procedures and openly released data and code, we provide a reproducible benchmark for comparing mosquito distribution and abundance models. This framework aligns with the growing use of artificial intelligence for modelling infectious-disease systems [26], while contributing specifically to ecological forecasting of invasive mosquito species and retaining direct relevance for vector surveillance and public-health preparedness.

The paper is organized as follows: The Materials and methods section describes the datasets, performance metrics, and technical details of our approach. We also introduce ***AI****edes*, our model composed of two modules: ***AI****edes*-Classifier for mosquito presence and ***AI****edes*-Counter for abundance prediction. The Results section presents model predictions across Europe and compares them with observed data. Final performance metrics are reported to support future benchmarking efforts. The Discussion section puts our approach, performance metrics and results in a wider modelling, infection control and public health context.

## 2. Materials and methods

### 2.1. Aedes albopictus datasets

*Mosquito presence prediction.* To model the presence distribution of *Ae. albopictus*, we used distribution maps [4] provided by the European Centre for Disease Prevention and Control (ECDC) and the European Food Safety Authority (EFSA) through their joint European network for medical and veterinary entomology (VectorNet) project, which collects highquality vector occurrence data from across Europe and the Mediterranean basin, including contributions from medical entomologists and public health experts [32]. The maps represent the distribution of mosquitoes at the NUTS3 administrative unit (AU) level, or equivalent. In Austria, Belgium, Denmark, Germany, The Netherlands, England (UK), Scotland (UK) and Wales (UK) NUTS2 was used instead. Administrative units are classified in 5 categories according to the presence of *Ae. albopictus*: i) established: where the species has been observed in at least one municipality within the AU and there is evidence of reproduction and overwintering, ii) introduced: where the species has been detected within the AU, but without confirmed establishment, iii) absent: where the species has not been detected within the AU despite surveillance efforts, iv) no data: where no sampling has been performed, and v) unknown: where it is not known if there are field studies on this species within the AU. For our analyses, we used data of type i) and iii) for confirmed presence and absence of mosquitoes respectively, relative to the year 2023, as the latest available when we started working on the model [33]. We did not consider data belonging to the other categories. Categories ii), iv) and v) do not provide a verified binary label: detection without confirmed establishment corresponds to neither established presence nor absence, while “no data” and “unknown” record the absence of surveillance rather than the absence of the species. Data pre-processing involved projecting the distribution data, provided at the AU level, onto a 12.5 km *×* 12.5 km grid covering European territory (see subsection 2.2.1). For each grid cell, we classify the presence or absence of *Ae. albopictus* following a masking procedure similar to that described by [18]. Specifically, all cells located within an AU where the species has not been detected (i.e. category iii of ECDC definition) were labelled with absence, while cells within an AU with records of the species were labelled with presence only if the local climate satisfied three predefined suitability criteria: annual precipitation *≥* 200 mm yr*^−^*^1^, mean temperature of the coldest month *≥ −*3 *^◦^*C, and annual mean temperature *≥* 10 *^◦^*C, corresponding respectively to the water requirement for larval development, to egg overwintering, and to the completion of development. Relative to [18], we applied a more conservative climatic envelope, taking the thresholds from the ecological limits reported in the ECDC guidelines [34]; they were fixed before any model fitting and were not tuned on the training or test data. The mask operates in one direction only: it can reassign an inherited presence label to absence, but never creates a presence record where surveillance reported none. The full definition is given in Section 1 of the Supporting Information. The resulting dataset comprises 45,724 grid cells. Before climatic filtering, 20,832 cells (45.6%) inherited a presence label from their administrative unit; the conservative mask retained 16,164 positive cells (35.4% of the final modelling dataset), so that 4,668 of the inherited positive labels (22.4%) were reassigned to absence. The data were partitioned into 80% for model development and 20% for final testing. The preprocessed, climate-paired versions of both datasets used in this work are described and publicly released in [33].

*Mosquito abundance prediction.* Given our objective to forecast mosquito abundance, the best data would be complete, high-resolution and geographically representative counts of adult female mosquitoes. In practice, however, such datasets are scarce and geographically heterogeneous, since comprehensive surveillance is costly and often limited to areas with higher incidence of Aedes-borne infections. Consequently, in accordance with established practices in insect ecology, we employed egg counts obtained through standardized ovitrap sampling protocols as a surrogate measure for adult female mosquito populations, since egg collection is easier and less costly than adult sampling, generally providing more extensive datasets[35, 36]. Accordingly, we used the AIMSurv dataset [37], the first pan-European harmonized surveillance effort for invasive Aedes mosquitoes and the most extensive source of standardized ovitrap-derived egg counts available in Europe. This dataset includes comprehensive mosquito surveillance data in the year 2020 across 24 countries, employing standardized methodologies to monitor the life cycle of invasive mosquitoes species belonging to the *Aedes* genus, including eggs, larvae, and adults. Sampling methods include harmonised procedures for frequency of collection, minimum length of sampling period and reporting, ovitrap positioning, larval sampling techniques, and adult trapping, covering extensive spatial and temporal scales to continuously monitor and record mosquito populations.

In the data pre-processing phase, we considered only the "egg" life stage of the data collected via ovitraps on *Ae. albopictus*. Since these records make up the majority of the AIMSurv dataset, their prevalence was likely to provide a sufficiently large sample for training and evaluating our machine learning model, increasing our confidence in the robustness of the results. Sampling frequency varied between 1 and 203 days, with around two-thirds of the data collected weekly or every two weeks. As our aim was to predict weekly egg-laying rates, we restricted our data to records with weekly sampling frequency, while allowing for some variability in the collection effort, including also 6 and 8 days. Using these selection criteria, we obtained a working dataset comprising 5,424 egg-stage records from 387 ovitraps. Around 80% of the readings contain zero eggs [33]. To allow direct comparison across samples collected at different intervals, we converted each egg count to a standardized weekly rate by dividing the observed count by the number of days in the sampling interval and multiplying by seven, thus expressing all values as eggs per 7-day period. Records reporting the same ovitrap and the same sampling interval were merged by averaging, which yields the modelling dataset of 5,129 weekly observations from 387 ovitraps. The resulting observations span 13 countries but are strongly unbalanced: Austria contributes 54.6% of the weekly observations yet only 0.1% of the non-zero ones, whereas Spain, Albania and Italy together provide 34.2% of the observations and 97.3% of the non-zero records.

To evaluate the generalization capability of our model beyond AIMSurv, we sought to test its performance on different data. We browsed the VectorNet (raw) dataset [38] for further suitable data. After applying the same selection criteria used for AIMSurv, i.e. focusing on the egg stage of *Ae. albopictus* and collection period of 6, 7 or 8 days, we identified suitable data only for the year 2021, consisting of 280 weekly egg collections from 40 ovitraps in the Lazio region of Italy, collected between 26 May and 16 November 2021, with up to 23 collections per ovitrap and uninterrupted weekly runs of at most 12 weeks, as described in [39].

### 2.2. Climatic and Biological Input Variables

Mosquito population dynamics are shaped by multiple factors, such as climate, photoperiod, and land use. Here, we restrict our analysis to climate variables —temperature, humidity, and precipitation— since they are key drivers [18, 36] and allow us to limit model complexity given the current size of our dataset. Other ecological and environmental variables could be integrated in future extensions of this work.

*Mosquito presence prediction.* For the classification task predicting the presence or absence of mosquitoes, we utilized climate data obtained from the European Union’s Earth observation program Copernicus [40], specifically the projections-cordex-domains-single-levels dataset. This dataset provides regional climate projections and historical simulations over Europe using a uniform grid with a horizontal resolution of approximately 0.11*^◦^*, corresponding to grid cells that are about 12.5 km wide.

To use a single climate source for the present analysis and for subsequent projections, we consistently employed the Copernicus dataset, which avoids discrepancies between historical reanalyses and future climate projections. For the presence/absence classification task, we considered the 12-month profiles of 2-m air temperature (*T*_12_*_m_*) and mean precipitation flux (*P*_12_*_m_*), calculated as monthly means over 2011–2020.

We used the 2011–2020 decadal monthly average of climatic variables to model the VectorNet *Ae. albopictus* distribution data from 2023, as the establishment of invasive mosquito species is typically shaped by broader climate patterns rather than year-to-year fluctuations. Decadal means provide a stable reference for both current and future projections, and the conditions in 2023 are expected to align closely with those in the early 2020s.

The climate fields were taken from the Global Climate Model and Regional Climate Model projections (GCM-RCM) for the medium-severity Representative Concentration Pathway (RCP4.5) [41], produced by the combined Max Planck Institute for Meteorology (MPI-M) and Swedish Meteorological and Hydrological Institute (SMHI) simulations. The same source therefore provides both the 2011–2020 decadal means used here and the projections required for future decades, which are not analysed in the present work.

We also used climate data from the WorldClim database [42] (1 km × 1 km resolution) for comparison with previous studies [18], with results reported in the Supporting Information (SI).

*Mosquito abundance prediction.* For the mosquito abundance predictions, we used, as climatic input variables, data from the the ERA5-Land dataset provided by the European Union’s Earth observation programme Copernicus [40]. ERA5-Land serves as a multidecadal, high-resolution reanalysis dataset that provides harmonized terrestrial environmental variables, offering enhanced 9-kilometer resolution numerical simulations of land surface processes [43].

We restricted the dataset to the European domain and extracted three key climatic variables: 2-meter air temperature, total precipitation, and 2-meter dew point temperature. For each sampling date, we extracted climate histories ending at the trap collection. We denote 2-m air temperature, total precipitation, and 2-m dew-point temperature by *T_W_*, *P_W_*, and *D_W_*, respectively, where *W* identifies the selected climate-history window. Climate inputs were represented either as daily summaries over 30, 60, or 90 days (30*d*, 60*d*, and 90*d*), or as weekly averages over the corresponding periods (5*w*, 9*w*, and 13*w*). Daily summaries are daily means for *T_W_* and *D_W_*, and daily total precipitation for *P_W_* . The daily sequences preserve shortto medium-term climatic variability, whereas weekly averaging provides a more compact representation with fewer input features.

We also evaluated previous weekly egg-laying rates, denoted by *E_W_*, with *W ∈ {*1*w,* 2*w}*. Models using measured previous rates receive the corresponding availability indicators *I_W_*, which allow the network to distinguish an observed zero value from an unavailable previous observation. The dedicated autoregressive model instead receives recursively generated previous predictions as *E_W_* ; no availability indicators are needed in this case, because the recursive sequence is defined at every step, taking the value zero before the corresponding lag exists. Climate-only configurations use the climate inputs alone. Models containing one or two previous weekly values were evaluated. Including recent egg-laying rates adds a temporal component that can represent the typical 7–10 day life cycle of *Ae. albopictus* from egg to adult [44] and the resulting density-dependent effects on egg-laying activity. This approach is consistent with previous findings that recent egg-laying observations can improve predictive accuracy by capturing information about recent reproduction [36].

Table 1 provides a comprehensive recap of all variables used in both classification and abundance prediction tasks.

**Table 1:** Input variables evaluated in the neural-network models. For the Counter, *W* identifies the selected history. The daily windows (30*d*, 60*d*, and 90*d*) contain 30, 60, and 90 daily climate summaries, respectively: daily means for *T_W_*and *D_W_*, and daily total precipitation for *P_W_* . The weekly windows (5*w*, 9*w*, and 13*w*) contain the corresponding numbers of weekly averages. Thus, the number in the subscript gives both the temporal window and the number of values supplied to the neural network for each climate variable. Climateonly configurations use the climate variables alone. Models using measured previous egg-laying rates receive *E_W_* together with *I_W_*, allowing the network to distinguish an observed previous rate from an unavailable value. In the dedicated autoregressive model, *E_W_* is instead supplied by the recursively generated previous prediction.

| **Task** | **Symbol** | **Description** |
|---|---|---|
| Classifier | *T*<sub>12*m*</sub> | Monthly mean 2-m air temperature for the 12 months, averaged over 2011–2020 |
|  | *P*<sub>12*m*</sub> | Monthly mean precipitation flux for the 12 months, averaged over 2011–2020 |
| Counter | *T*<sub>*W*</sub> | 2-m air temperature; *W* ∈ {30*d*, 60*d*, 90*d*, 5*w*, 9*w*, 13*w*} |
|  | *P*<sub>*W*</sub> | Total precipitation; *W* ∈ {30*d*, 60*d*, 90*d*, 5*w*, 9*w*, 13*w*} |
|  | *D*<sub>*W*</sub> | 2-m dew-point temperature; *W* ∈ {30*d*, 60*d*, 90*d*, 5*w*, 9*w*, 13*w*} |
|  | *E*<sub>*W*</sub> | Previous weekly egg-laying rate; *W* ∈ {1*w*, 2*w*} |
|  | *I*<sub>*W*</sub> | Availability indicator for *E*<sub>*W*</sub>; *W* ∈ {1*w*, 2*w*} |

### 2.3 Performance Metrics

#### 2.3.1. Presence prediction task

To evaluate the ability of our models to predict the presence or absence of *Ae. albopictus* in Europe, and to provide a basis for comparison with other approaches, we used four standard evaluation metrics, all computed on held-out data:

• **Accuracy:** Proportion of correctly predicted locations:

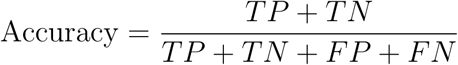

• **Sensitivity (True Positive Ratio):** Proportion of presence locations correctly identified:

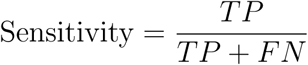

• **Specificity (True Negative Ratio):** Proportion of absence locations correctly identified:

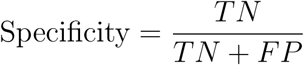

**Critical Success Index (CSI):** Proportion of correctly predicted presence locations relative to all locations where presence was either observed or predicted:

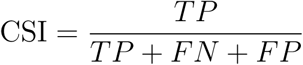

Here, *TP*, *TN*, *FP*, and *FN* denote the number of locations where the species is present and predicted present (true positives), absent and predicted absent (true negatives), absent but predicted present (false positives), and present but predicted absent (false negatives), respectively. These metrics provide a comprehensive assessment of the model’s performance in distinguishing between presence and absence cases.

#### 2.3.2. Abundance prediction task

For the abundance prediction task, we evaluated model performance on the observations of traps excluded from the corresponding training sets, to measure the ability of the model to generalize to ovitraps not seen during training. We used the coefficient of determination (*R*^2^), which measures how well the predicted values match the observed weekly egg counts. It is defined as:

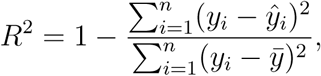

where *y_i_* are the observed values, *ŷ_i_* are the predicted values, and *ȳ* is the mean of the observed values.

We also computed the root mean squared error (RMSE), which gives the average size of the prediction errors:

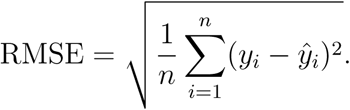

Because the weekly rates are strongly right-skewed, we additionally report the coefficient of determination on the logarithmic scale, obtained by applying the log(1+*x*) transformation to the observed and predicted values before evaluating the same expression:

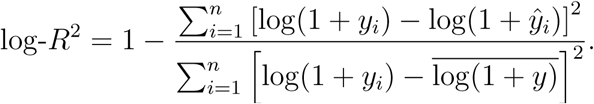

This measure weights proportional rather than absolute deviations, so that agreement at low and intermediate rates is not dominated by the largest observations.

Finally, to check whether the model can correctly distinguish between the absence and presence of mosquito eggs, we calculated the accuracy in detecting zero versus non-zero values. This was done by converting both observed and predicted egg counts into binary values (zero or non-zero), and computing the proportion of correct predictions. An observation counts as non-zero when at least one egg was recorded, and a prediction counts as non-zero when the predicted weekly rate reaches one egg, since the model outputs continuous positive values.

### 2.4. Neural Network Models

***AI****edes* is a neural network-based model comprising two components: ***AI****edes*-Classifier and ***AI****edes*-Counter. The first estimates the probability of presence of *Ae. albopictus*, while the second predicts the ovitrap-derived weekly egg-laying rate, which can serve as a general proxy for local *Ae. albopictus* abundance. They are implemented in Python using the PyTorch framework [45]; the code and the final neural network model are freely accessible in [46]; the curated datasets are described and released in [33]. For the Classifier, the data were randomly divided into a model-development set representing 80% of the available data and a test set consisting of the remaining 20%. Model configurations were compared by five-fold validation within the model-development set. For the Counter, complete traps were divided into five groups; each group served in turn as the test set, while the observations from the other four groups were used for training.

Figure 1 shows the analysis workflow used by both components to estimate the probability of presence and weekly egg-laying rate of *Ae. albopictus* from climate and biological variables at a given location. The neural-network drawings are schematic; their arrows show the feed-forward direction of information flow, whereas task-specific architectures were selected through validation.

**Figure 1:**
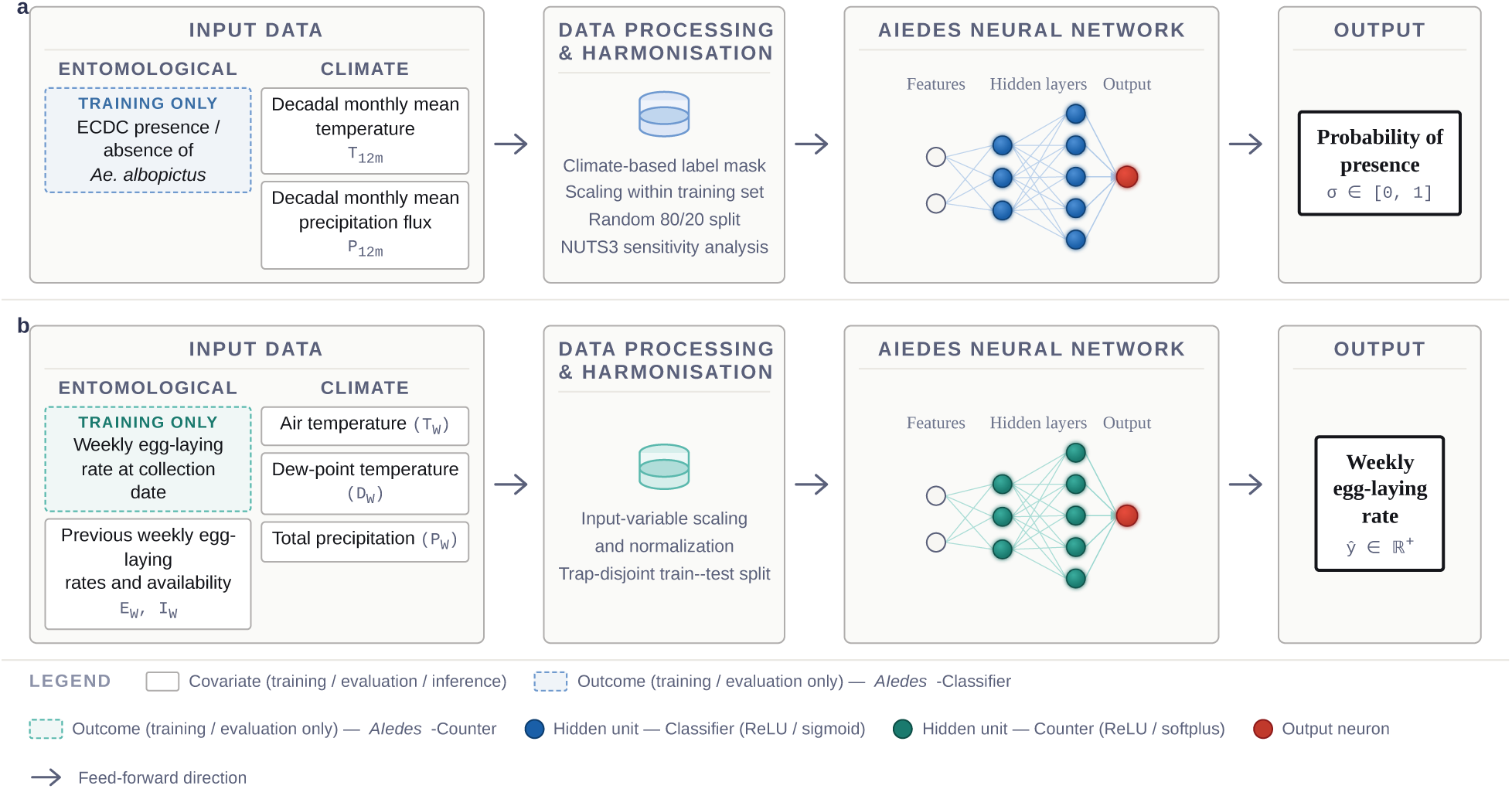
Schematic representation of the analysis workflow for the two components of ***AI****edes*. **(a) *AI****edes*Classifier: the model uses the 12-month profiles of 2-m air temperature (*T*_12_*_m_*) and precipitation flux (*P*_12_*_m_*) to predict the probability of presence of *Ae. albopictus*. Climate-based label masking is applied before a random 80/20 train–test split, whereas scaling parameters are estimated using the training data only within each fit; NUTS3-level splitting provides an additional sensitivity analysis. **(b) *AI****edes*-Counter: the model uses climate histories of air temperature (*T_W_* ), dew-point temperature (*D_W_* ), and total precipitation (*P_W_* ) to predict the weekly egg-laying rate *ŷ* *∈* ℝ^+^. Depending on the configuration, it can also use previous weekly egg-laying rates (*E_W_* ) and their availability indicators (*I_W_* ). In the dedicated autoregressive configuration, previous egg-laying rates are recursively generated predictions. Here, *W* denotes the selected history window, as defined in Table 1. The Counter workflow uses trap-disjoint validation and, for final models, five random initializations per fold.

#### 2.4.1. AI edes-Classifier

The goal of the ***AI****edes*-Classifier module is to predict the probability of presence of *Ae. albopictus* across Europe in a given calendar year. The input variables consist of monthly climatic measurements, aggregated from the original hourly data: temperature and precipitation. The model outputs a value between 0 and 1, which represents the probability of *Ae. albopictus* being present at a given location during a calendar year. By convention, a predicted probability greater than 0.5 is considered indicative of the presence of *Ae. albopictus*, while a probability of 0.5 or lower is interpreted as absence.

The hidden layers use the ReLU (Rectified Linear Unit) activation function, while a sigmoid transformation of the final output gives probabilities in the [0, 1] range. When no hidden layers are present, the model is mathematically equivalent to logistic regression. Within the 80% model-development partition, we compared temperature-only and temperature-plusprecipitation inputs and network architectures containing zero to three hidden layers by five-fold stratified validation. The configuration with the highest mean validation accuracy was then evaluated on the untouched 20% test partition using five independent random initializations whose probabilities were averaged. Input scaling was estimated using only the training data in each fit. The model was trained using the binary cross-entropy loss function [47] and optimized with the Adam optimizer [48], with a learning rate of 0.002. Classifier models were trained for a fixed 500 epochs, with the training and validation losses recorded at every epoch. The fixed schedule was used for all classifier fits, without selecting an intermediate epoch.

For the full-coverage comparison in Figure 4, the selected configuration and the nohidden-layer reference were evaluated by five-fold stratified out-of-fold prediction, repeated with five random initializations per fold. Each cell was predicted only by models trained without that cell, and the five probabilities were averaged. As a spatial-transfer sensitivity analysis, we repeated this comparison using five-fold stratified grouped validation with complete NUTS3 regions as groups, so that no region occurred in both training and held-out data within a fold.

Figure 2a illustrates the typical behavior of the training and validation loss functions over epochs. To optimize model performance, we conducted a series of tests varying both the number of hidden layers and the number of units per layer (Figure 2b). Architectures ranged from no hidden layers (equivalent to logistic regression) to configurations with up to three hidden layers containing 50 units each. We evaluated model performance using the accuracy metric, considering two combinations of input variables: (i) monthly averaged temperature only, and (ii) monthly averaged temperature together with monthly total precipitation.

**Figure 2:**
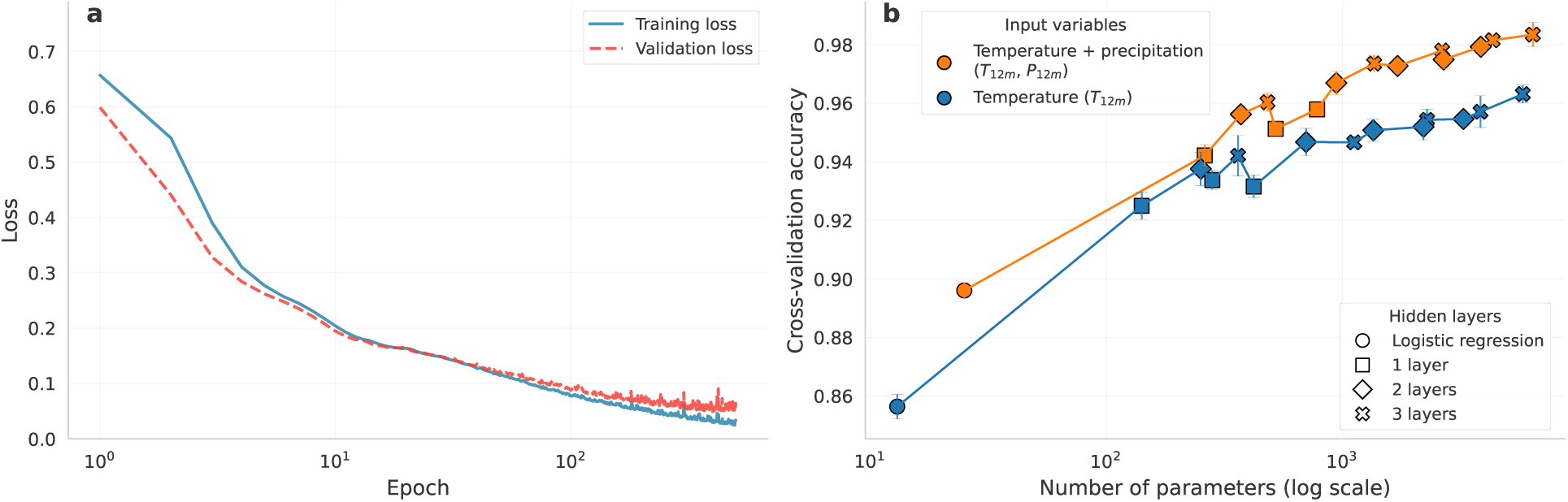
Training dynamics and model selection of *AIedes*-Classifier. **(a)** Representative training and validation loss curves for the selected model using temperature and total precipitation and three hidden layers with 50 units each ([50, 50, 50]). The curves show one validation fold within the 80% modeldevelopment partition. **(b)** Mean validation accuracy for temperature only and temperature plus precipitation as a function of the total number of model parameters (log scale). Points represent architectures ranging from no hidden layer, equivalent to logistic regression, to three hidden layers; error bars show the standard deviation across the five validation folds.

The results, shown in Figure 2b, indicate that increasing the number of layers and hidden units (and thus the total number of parameters) continues to yield performance gains, even at the highest tested values. However, the improvements become increasingly marginal and, as illustrated by the log-scaled x-axis, these small gains require exponentially larger increases in the number of parameters.

The inclusion of both monthly average temperature and total precipitation significantly improved the predictive accuracy, leading to an accuracy value close to 99% for the model featuring the highest number of hidden layers and parameters. This is therefore the final model configuration chosen for testing (see Results section).

#### 2.4.2. AI edes-Counter

The ***AI****edes*-Counter module is designed to predict the weekly egg-laying rate of *Ae. albopictus* collected using ovitraps, as defined in the AIMSurv project, using climate variables from previous periods at each specific location.

The input variables consist of the climatic and biological factors described in the Data section. Model performance was evaluated using five-fold grouped cross-validation by trap, a standard procedure in which complete groups are withheld in turn. The traps were divided into five groups, and all observations from one group formed the evaluation set while the remaining four groups formed the training set. Consequently, every evaluation observation came from a trap that was excluded from the corresponding training data. Climate and mosquito-history inputs were scaled using statistics estimated from the training traps in each fold.

The selected Counter uses a Weighted Mean Squared Logarithmic Error (WMSLE) loss function to accommodate the strongly right-skewed distribution, in which approximately 80% of observations are zero. Weighted Mean Squared Error (WMSE) and a zero-inflatednegative-binomial-inspired loss were also evaluated, with the complete comparison reported in the Supporting Information.

We used feed-forward neural networks with ReLU hidden-layer activations and a final softplus activation for the predicted weekly rate. The tested hidden-layer architectures were [32], [64], [32, 16], and [64, 32], combined with dropout rates of 0, 0.2, and 0.4 [49]. The initial sensitivity screen trained every configuration once in each of the five trap folds. Figure 3 reports the mean held-out-trap *R*^2^ and its standard error across these folds, retaining every completed training run. Configurations selected for final evaluation were subsequently trained from five independent random initializations in each fold, yielding 25 retained models for each selected configuration.

Figure 3 shows that daily climate inputs generally provide higher held-out-trap *R*^2^ than weekly averages of the same 90-day history. Adding the measured oneand two-week egglaying rates generally improves performance, while differences among network architectures and dropout rates are smaller and vary across trap folds. The selected observed-history configuration combines *T*_90_*_d_*, *P*_90_*_d_*, *D*_90_*_d_*, *E*_1_*_w_*, and *E*_2_*_w_*, with availability indicators *I*_1_*_w_* and *I*_2_*_w_*, with a [64] hidden layer and dropout of 0.2. The broader comparisons across climatehistory lengths and loss functions are presented in the Supporting Information.

**Figure 3:**
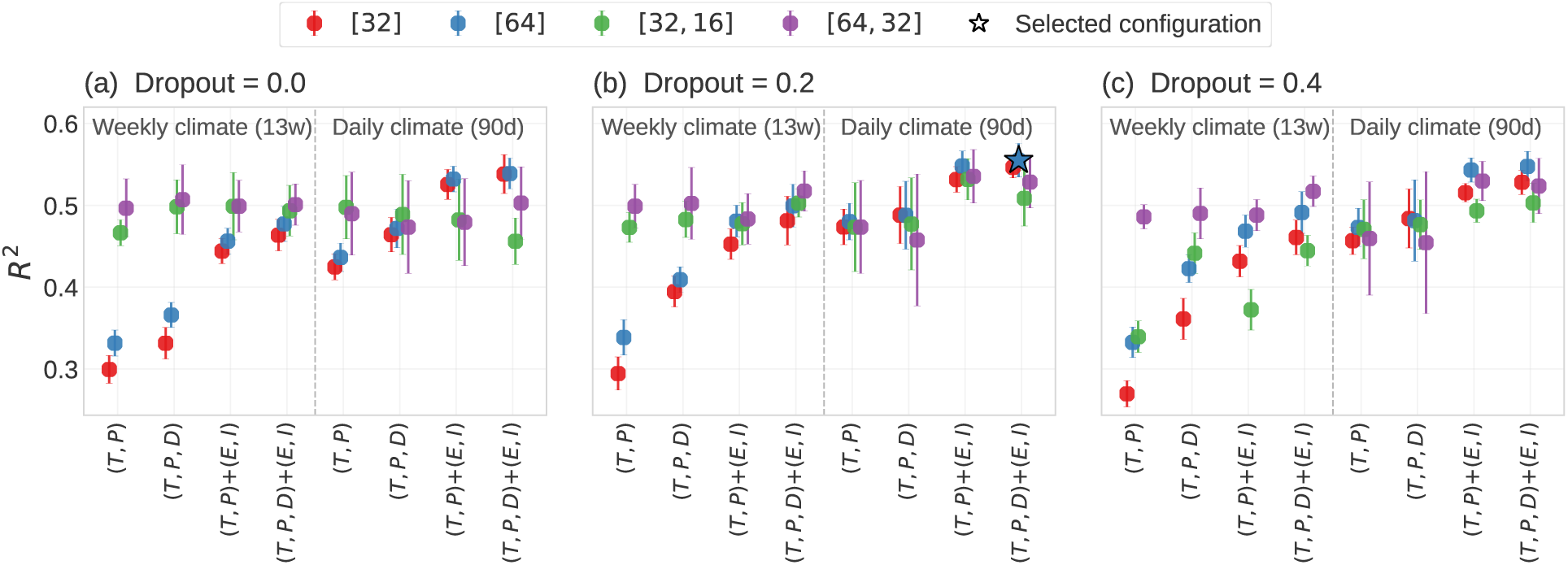
Counter input and architecture sensitivity. Mean held-out-trap *R*^2^ for WMSLE-trained models using 90-day climate histories represented as 90 daily values or 13 weekly averages. Within each temporal representation, the x axis shows four input sets: *T, P* ; *T, P, D*; *T, P* with *E, I*; and *T, P, D* with *E, I*, where *E, I* denotes *E*_1_*_w_*, *E*_2_*_w_*, *I*_1_*_w_*, and *I*_2_*_w_*. Panels show dropout rates of 0.0, 0.2, and 0.4, with the dashed line separating weekly and daily climate representations. Points identify the four tested hidden-layer architectures, and error bars show the standard error across the five trap-disjoint folds. Each configuration was screened using one training seed per fold, and every completed run was retained. The star identifies the observed-history configuration selected for subsequent evaluation; this configuration was then trained using five independent seeds in each fold. Comparisons of 30-, 60-, and 90-day histories and alternative loss functions are reported in the Supporting Information.

Table 2 compares prediction using measured previous egg-laying rates with two autonomous alternatives on a common cohort. Recursive prediction requires an uninterrupted daily climate series for each trap, which the archived model inputs provide for 375 of the 387 traps; the remaining 12 traps, contributing 93 observations, retain gaps between the stored 90-day windows and are therefore excluded, so that all four modes are evaluated on the same 5,036 observations. The observed-history and climate-only configurations were selected independently from their respective WMSLE sensitivity analyses. The dedicated autoregressive architecture was selected through a focused comparison between [64] and [64, 32] networks. Measured previous values provide the highest predictive performance when continuing trap observations are available. Applying that model recursively reduces *R*^2^ from 0.565 to 0.456, whereas training directly with recursively generated previous values increases autonomous-forecast *R*^2^ to 0.535. The dedicated autoregressive model has log-*R*^2^ and accuracy comparable to the climate-only configuration, together with higher original-scale *R*^2^ and lower RMSE. We therefore use the dedicated autoregressive configuration for projections that proceed without continuing trap observations.

**Table 2:** Comparison of Counter prediction modes on the common cohort of 5,036 observations from 375 traps. The observed-history model uses the daily climate histories *T*_90_*_d_*, *P*_90_*_d_*, and *D*_90_*_d_*, together with the measured previous egg-laying rates *E*_1_*_w_* and *E*_2_*_w_* and availability indicators *I*_1_*_w_* and *I*_2_*_w_*, and has a [64] network with dropout 0.2. The dedicated autoregressive model uses the same climate inputs with recursively generated *E*_1_*_w_* and *E*_2_*_w_*, no availability indicators, and a [64, 32] network with dropout 0.2; the focused exploration of autoregressive architectures is reported in the Supporting Information. The climate-only model is the best-performing configuration among the WMSLE climate-only models, selected by mean heldout-trap *R*^2^; it uses *T*_13_*_w_*, *P*_13_*_w_*, and *D*_13_*_w_*, a [64, 32] network, and dropout 0.0. Log-*R*^2^ uses the log(1 + *x*) transformation.

| Mode | History: training / evaluation | Inputs | *R*<sup>2</sup> | log-*R*<sup>2</sup> | RMSE | Acc. |
|---|---|---|---|---|---|---|
| Observed history | Measured / measured | (*T*, *P*, *D*)<sub>90*d*</sub>, (*E*, *I*)<sub>1*w*,2*w*</sub> | 0.565 | 0.678 | 26.36 | 0.921 |
| Observed-history model used recursively | Measured / predicted | (*T*, *P*, *D*)<sub>90*d*</sub>, (*E*, *I*)<sub>1*w*,2*w*</sub> | 0.456 | 0.649 | 29.48 | 0.916 |
| Dedicated autoregressive | Predicted / predicted | (*T*, *P*, *D*)<sub>90*d*</sub>, *E*<sub>1*w*,2*w*</sub> | 0.535 | 0.663 | 27.25 | 0.919 |
| Climate only | – | (*T*, *P*, *D*)<sub>13*w*</sub> | 0.519 | 0.664 | 27.72 | 0.921 |

## 3. Results

### 3.1. Predicting Local Mosquito Presence and Absence

Here we present the results of the ***AI****edes*-Classifier model, trained on presence and absence data. For this analysis, the selected input variables are monthly average temperature and total precipitation, as described in Methods. Figure 4 compares predictions from the simplest neural network tested a model with no hidden layers, statistically equivalent to logistic regression (Figure 4a) and those from the best-performing architecture, consisting of three hidden layers with 50 units each (Figure 4b). These models predict the geographical distribution of *Ae. albopictus* using observational data (see Data section). The input variables are monthly decadal averages of temperature and total precipitation values over the decade 2011–2020, derived from the Copernicus dataset.

The output of these models is the probability of presence for *Ae. albopictus*, where probabilities above 0.5 are interpreted to indicate predicted presence and probabilities below 0.5, to indicate absence. On the untouched 20% test partition, the five-model ensemble achieved an accuracy of 0.990, sensitivity of 0.982, specificity of 0.995, and CSI of 0.973. The full-coverage random out-of-fold comparison shown in Figure 4 gave closely corresponding values: accuracy 0.991, sensitivity 0.988, specificity 0.992, and CSI 0.974 for the selected neural network, compared with 0.895, 0.853, 0.918, and 0.743, respectively, for the no-hiddenlayer logistic reference. The nonlinear network therefore provided a substantial improvement over the simpler reference model. To assess geographic transfer, we additionally performed the same comparison while withholding complete NUTS3 regions. Each observation was evaluated only by models trained without data from its NUTS3 region. Across the five held-out regional folds, the selected network had mean accuracy 0.920 *±* 0.007, sensitivity 0.890 *±* 0.048, specificity 0.936 *±* 0.025, and CSI 0.797 *±* 0.020. The corresponding logistic-reference values were 0.888 *±* 0.020, 0.844 *±* 0.035, 0.911 *±* 0.038, and 0.728 *±* 0.036. Absolute performance was lower and more variable than in the random-split evaluation, while the selected neural network maintained its improvement over the logistic reference for all four metrics. We therefore interpret the NUTS3 analysis as an additional regional-transfer stress test rather than as the principal performance estimate (see the SI for more information).

### 3.2. Abundance Predictions

Figure 5 compares observed weekly egg-laying rates with predictions from the selected observed-history configuration. The training predictions achieved *R*^2^ = 0.712 and log-*R*^2^ = 0.850, while predictions for traps excluded from model training achieved *R*^2^ = 0.566, log*R*^2^ = 0.680, and RMSE = 26.12 across 5,129 observations from 387 traps. Weekly egg-laying rates are strongly right-skewed, and their absolute fluctuations tend to increase with the magnitude of the observations because of stochastic oviposition and local sampling variability.

**Figure 4:**
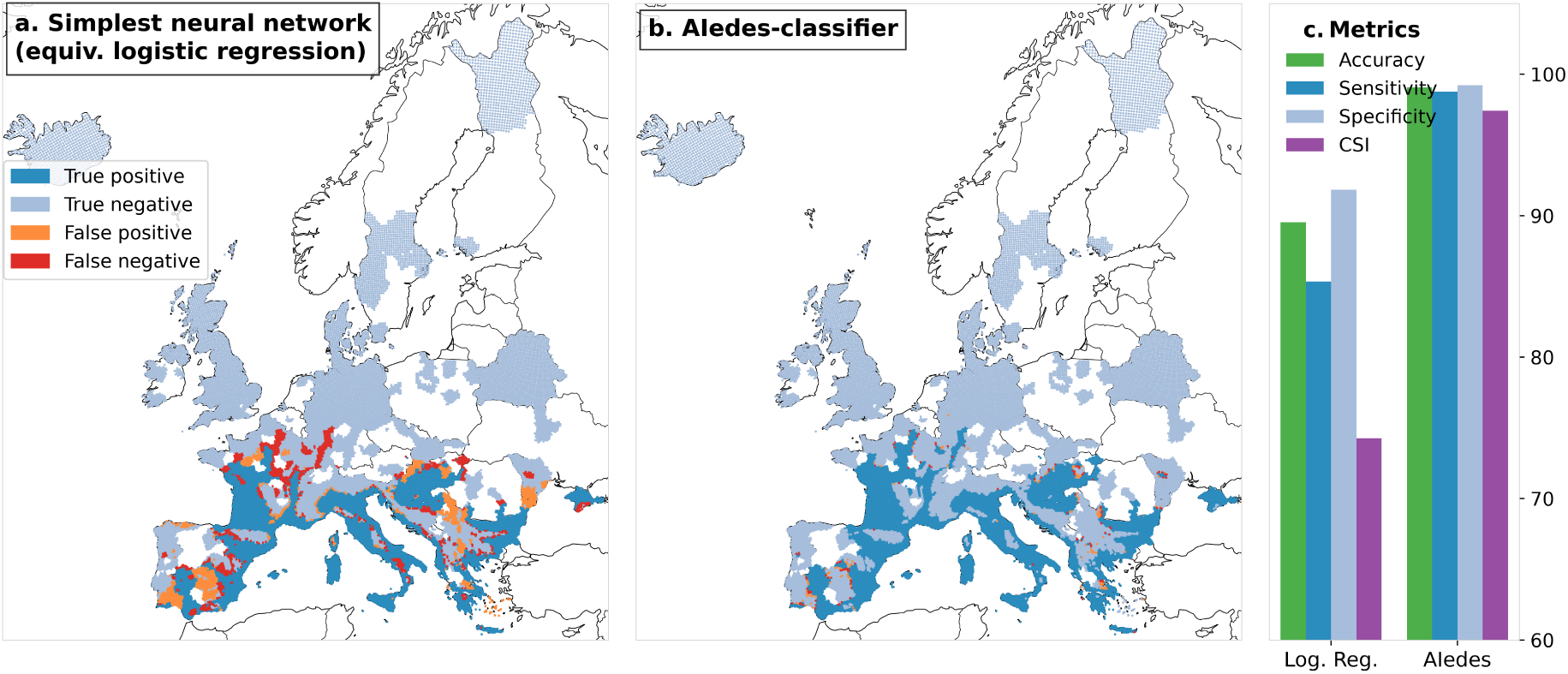
Random out-of-fold classification of *Ae. albopictus* presence and absence. Each grid cell was classified using the mean probability from five models that were not trained on that cell. a. Results from the simplest architecture, featuring no hidden layers and equivalent to logistic regression. b. Results from the selected model, featuring three hidden layers with 50 units each. c. Pooled accuracy, sensitivity, specificity and critical success index (CSI) across all held-out predictions.

**Figure 5:**
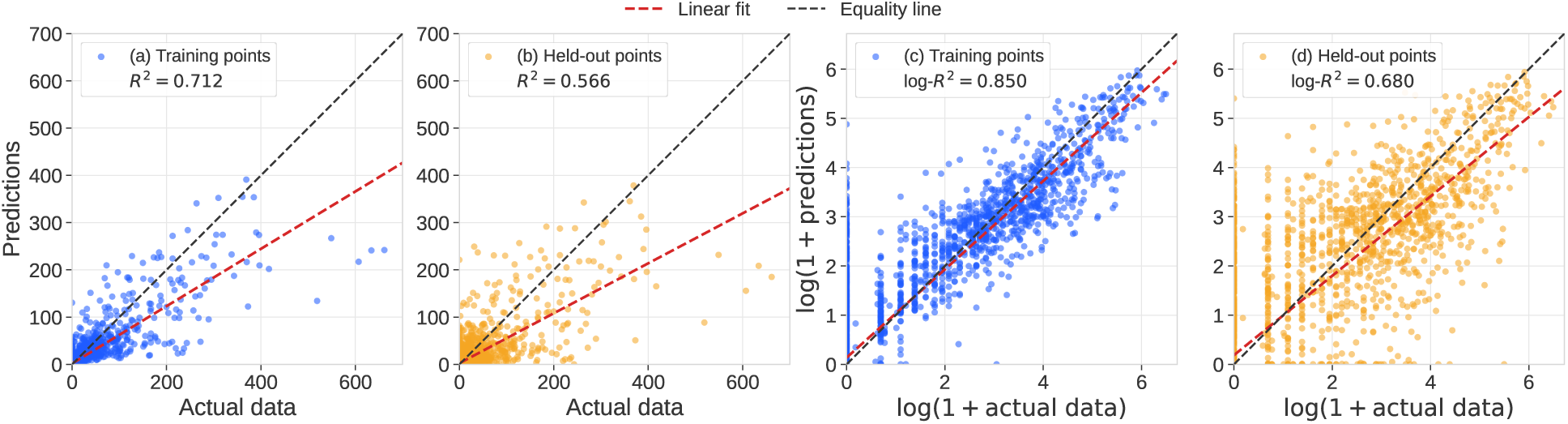
Observed and predicted weekly egg-laying rates. Panels (a) and (b) show the training and heldout-trap predictions on the original scale, respectively; panels (c) and (d) show the corresponding results after log(1 + *x*) transformation. Red lines show the linear fits and black dashed lines indicate equality. The held-out predictions are means from the five models that excluded the corresponding trap from training. Predictions comprise 5,129 observations from 387 traps. Points lie slightly below the equality line at the highest rates, while the mean difference between predicted and observed rates remains small, indicating no overall systematic bias (see Section 4 in the SI).

The log(1 + *x*) scale therefore complements the original-scale assessment by reducing the influence of occasional high values and showing proportional agreement more clearly across low and intermediate rates.

The same ensemble was also applied to the independent ECDC/EFSA VectorNet collections from Lazio, which are reported separately from the AIMSurv results. Across the 280 collections the observed-history configuration obtains *R*^2^ = *−*0.004, log-*R*^2^ = 0.170 and RMSE = 69.8, with a zero/non-zero accuracy of 0.754 against the 0.736 expected from assigning every week to the non-zero class. In a post hoc split of the season, performance is better in the first part than in the second: until the end of August it reaches *R*^2^ = 0.072, log*R*^2^ = 0.360 and accuracy 0.821 on 162 collections, while from September onward it reaches *R*^2^ = *−*0.071, log-*R*^2^ = *−*0.076 and accuracy 0.661 on 118 collections. In both periods the agreement remains well below the values obtained on the pan-European AIMSurv data.

Figure 6 displays weekly egg-laying rates from selected traps, comparing observations with the corresponding ***AI****edes*-Counter predictions. The figure presents nine AIMSurv time series and three ECDC/EFSA VectorNet time series. The AIMSurv traps were selected by ranking the filtered weekly records within each country by their number of observations, retaining three series from Albania and two each from Austria, Italy, and Spain; ties were resolved by trap identifier. The three VectorNet series, all from Italy in 2021, were selected independently by the same criterion. Within each subplot, the points show weekly observations and ensemble-mean predictions, and the vertical bars show their prediction variability. Centred three-week moving averages facilitate comparison of temporal trends.

**Figure 6:**
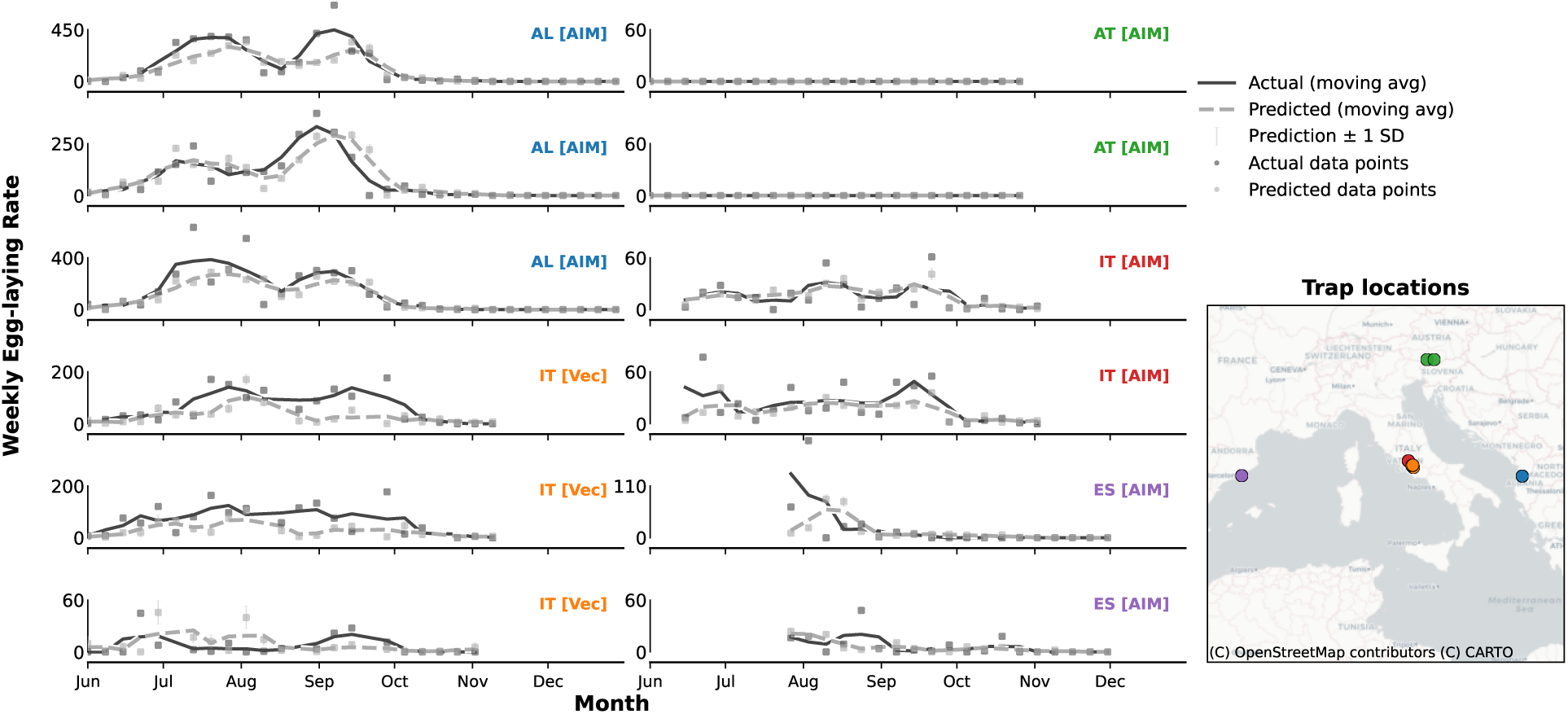
Observed and predicted weekly egg-laying rates at selected AIMSurv and VectorNet traps. Within each displayed AIMSurv country, traps were ranked by the number of eligible weekly observations after filtering, and the longest three series from Albania and the longest two from Austria, Italy, and Spain were retained. The three VectorNet traps were selected independently using the same criterion. Points show observations and ensemble-mean predictions. Vertical bars show *±*1 standard deviation across the five models used for AIMSurv predictions and across the 25-model ensemble used for VectorNet predictions. Solid and dashed curves show centred three-week moving averages of observations and predictions, respectively. The y axes differ among panels.

Finally, Figure 7 presents monthly averages of the predicted weekly egg-laying rates for June through September 2020. At each grid cell, the dedicated autoregressive ensemble combines the 90-day daily climate history with its own predictions from 7 and 14 days earlier. The recursive sequence is initialized with zero previous egg-laying values and is continued through the calendar year. The maps show the ensemble mean across the 25 models and describe the spatial and seasonal pattern produced by this model configuration.

**Figure 7:**
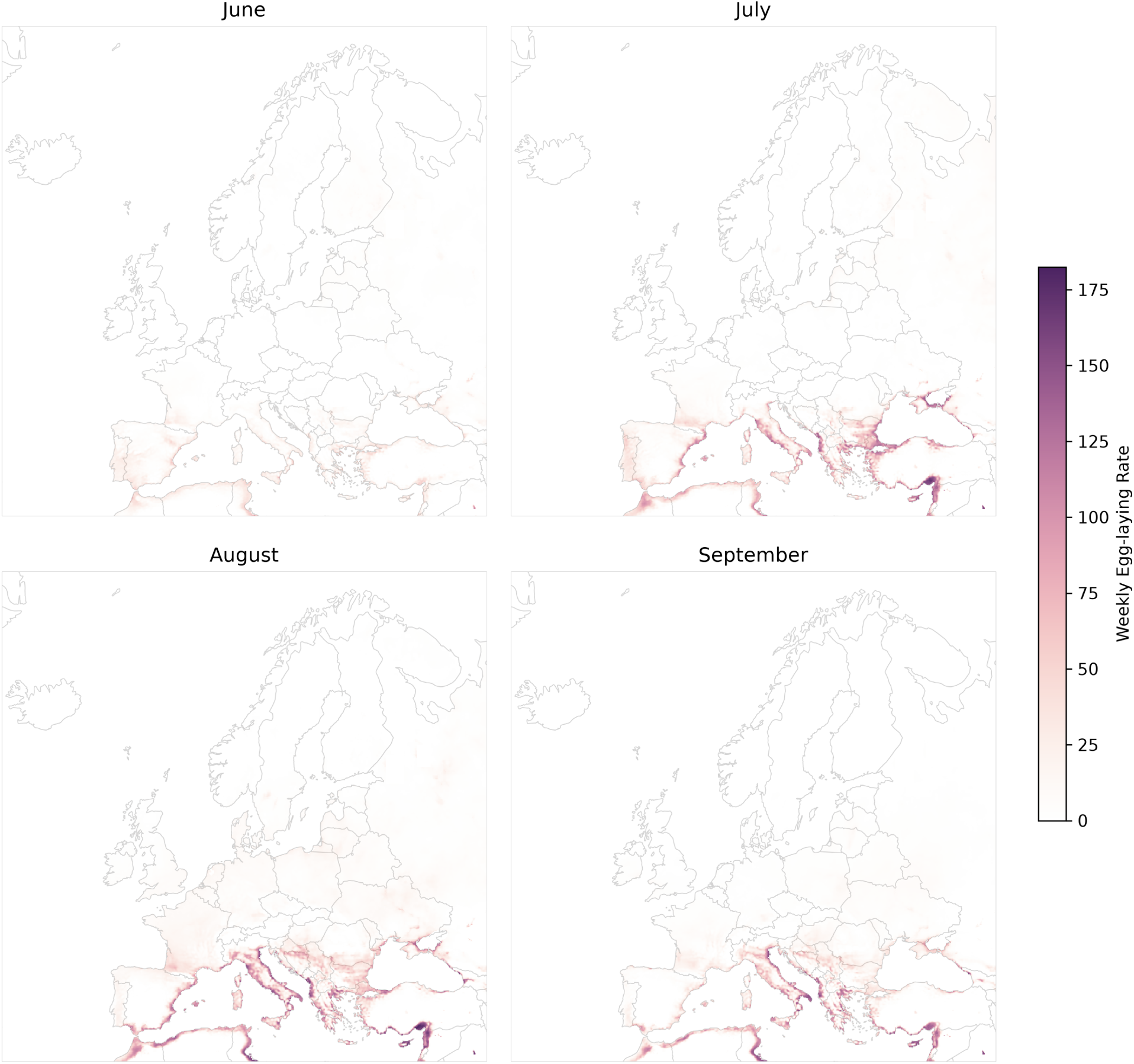
Monthly mean predicted weekly egg-laying rates of *Ae. albopictus* across Europe from June to September 2020. Predictions were generated by the dedicated autoregressive Counter ensemble, comprising 25 models trained across five trap-based folds and five random initializations. The 7and 14-day previous egg-laying inputs were recursively generated during both training and projection; the sequence was initialized with zero values before the respective lag became available. Mapped values are ensemble means.

## 4. Discussion

The field of modeling vector mosquitoes’ spatial distribution and abundance has long faced considerable challenges and limitations in developing comparable and reproducible approaches. Models are often applied to data on small spatial extents [9, 10, 13, 14, 25, 30], rely on data sets and quantities that are not easily reproducible [11], or feature only qualitative analyses without robust performance metrics [15, 18]. These limitations have made systematic comparisons across models difficult, especially for predicting vector abundance, where data are typically more scarce, heterogeneous, and collected under different protocols, not necessarily for the purpose of predicting vector abundance. In addition, models often predict non-comparable quantities and use code that is either unavailable or not user-friendly.

Recent developments in data availability and methodology have created new opportunities to improve the quality and reliability of mosquito vector distribution estimation. Thanks to efforts such as the ECDC mosquito presence maps and the AIMSurv European collaboration, large-scale datasets are now available, enabling among other things the application of novel and powerful machine learning approaches. Building on these advances, here we present ***AI****edes*, a new, fully data-driven, validated and replicable approach for estimating mosquito presence and abundance using neural networks.

By combining the well-curated datasets from ECDC/EFSA VectorNet [34], the AIMSurv project [37], and Copernicus climate data, we establish a framework that describes mosquito distributions on a continental scale and also applies rigorous validation and performance assessment via formal accuracy and *R*^2^ statistics. This framework enables transparent evaluation and facilitates future comparisons with other modeling approaches using shared or comparable datasets.

Our ***AI****edes*-Classifier reproduced the presence–absence labels with an accuracy of 0.991 in the full-coverage out-of-fold evaluation, compared with 0.895 for the logistic formulation applied to the same inputs, which is the functional form adopted in comparable distribution models [18]. Sensitivity and specificity in the same evaluation are 0.988 and 0.992, respectively (Figure 4). The ***AI****edes*-Counter reproduced weekly egg-laying rates at traps excluded from training with *R*^2^ = 0.566 and log-*R*^2^ = 0.680.

Withholding complete NUTS3 regions from training reduces accuracy to 0.920 *±* 0.007 and CSI to 0.797 *±* 0.020, compared with 0.888 *±* 0.020 and 0.728 *±* 0.036 for the logistic reference. This reduction may reflect the removal of part of the climatic range from the training data when entire regions are withheld, which could require predictions for conditions that are sparsely represented during training. The advantage over the logistic reference is retained under both validation schemes.

***AI****edes* Classifier is able to identify correctly both the overwhelming majority of grid cells where *Ae. albopictus* is present and at the same time is not prone to incorrectly assigning presence in locations where the insect is absent.

Our best model limits areas with misclassifications to parts of Hungary, Serbia and Central Spain. A comparison with our map of estimated egg-laying rates (Figure 7) indicates that these areas feature very low expected number of mosquitoes, suggesting that the practical implications of such misclassification by the presence model would in fact be minimal.

For the Counter model, the ensemble predictions for traps excluded from training achieved *R*^2^ = 0.566, log-*R*^2^ = 0.680, and RMSE = 26.12 (Figure 5). The grouped evaluation therefore measures performance at surveillance traps that were not used to train the corresponding models. Performance is higher on the logarithmic scale, reflecting the broad and right-skewed distribution of ovitrap-derived weekly egg-laying rates and the greater dispersion among the highest observations. The remaining variation is consistent with the strong short-term and local fluctuations of mosquito population dynamics and egg-laying activity. Trap placement, competing breeding sites, sampling conditions, and micro-environmental variation can all affect observed egg-laying rates at scales finer than the climatic inputs. Longer time series, replicated observations, and higher-resolution environmental information may therefore improve predictive accuracy.

Figure 6 illustrates predictions for the country-balanced longest series selected according to the number of eligible weekly observations. In the three VectorNet time series, the model does not capture the secondary peak in weekly egg counts between September and October. The underlying reason is difficult to assess. The selected VectorNet time series were collected in a separate elevation-focused study. The available metadata do not establish that all aspects of trap placement, collection, and processing were equivalent to the AIMSurv protocol. These time series may therefore reflect environments and procedures not represented in the AIMSurv dataset, and the relationships between input and output variables in this context may differ from the one captured during ***AI****edes* training on AIMSurv data. Specifically, given the high altitude and location of the sampling sites, it is plausible that local microclimatic conditions, especially in valleys or high-plateau environments, may not have been adequately represented in the training data set or sufficiently resolved in the climatic grid, which may have influenced the egg-laying-rate predictions.

Our predicted *Ae. albopictus* egg laying rates for throughout Europe (Figure 7) exhibit a notable cline of increase of egg-laying rates towards southern Europe, a pattern consistent with climatic conditions in Europe’s subtropical belts, much more suitable for breeding of the species. We also observe considerably higher predicted egg-laying rates in the coastal areas of the Mediterranean, but also of the Black and Azov Seas further to the north. The four panels of Figure 7 also show that mosquito abundance exhibits a marked summer seasonal pattern with a peak during the months of July and August the period of the year with higher day and night temperatures in combination with long periods of daylight. European countries positioned further north tend to peak in vector abundances later in the summer season, in August and even in September, as the most favourable weather conditions are likely reached later in these locations.

In order for ***AI****edes*-Classifier model to have public health utility, the next step would be to go beyond reproducing the current state of knowledge of *Ae. albopictus* distribution status, and make predictions for those areas where *Ae. albopictus* is expected to be present in the absence of current surveillance data (e.g. in areas currently marked as ’no data’ or ’unknown’ in the map), so that surveillance can be targeted to those areas. A further step might even be to try to make distribution predictions for the future depending on climate change scenarios. However, life-history and ecological considerations will need to be taken into account when interpreting such maps of predicted *Ae. albopictus* distribution in Europe: In the course of its migration to, and establishment in new areas in Europe the species is unlikely to have yet fully occupied its ecological niche across the continent. For instance, in the Netherlands, the species is not yet established despite suitable climatic conditions, due to active elimination efforts in the country. We excluded this region from our training dataset, however, similar situations may exist in locations where *Ae. albopictus* is still in the process of expansion into new climatically suitable areas. Such locations are marked as negative not because of unsuitable conditions but because the invasion process is still incomplete. The species could therefore spread further under current climate conditions, potentially limiting the accuracy of predictions based on present-day distribution data. Because the ECDC categories are assigned from surveillance evidence at the administrative-unit level, the information content of an absence label depends on the surveillance carried out in that unit; our grid-level labels inherit this structure, so a negative label denotes the absence of detection rather than demonstrated unsuitability. A further methodological consideration concerns the construction of grid-level presence–absence labels. Our downscaling procedure combines NUTS3-level inputs with spatially structured Copernicus climate predictors, so neighbouring grid cells can share similar covariates and the same aggregated label. Because the classifier uses climate predictors, it can also learn the broad climatic limits encoded by the suitability mask; the reported metrics therefore quantify agreement with a climatefiltered distribution target rather than validation against independent cell-level occurrence records, which are not available at the European scale. The random-split evaluation measures interpolation within the sampled European geographic and climatic domain, which is the principal prediction setting considered here. The additional NUTS3 analysis addresses the distinct question of transfer to administrative regions excluded from training. It yields lower and more variable absolute performance, while preserving the improvement of the selected neural network over the logistic reference (see Supporting Information). Because NUTS3 regions are administrative rather than climatic units, this analysis characterizes geographic transfer rather than extrapolation to entirely unseen climatic conditions.

Similarly, for the ***AI****edes* Counter model, several considerations will need to be taken into account when interpreting predicted *Ae. albopictus* population dynamics time series in Europe. *First*, the training data originate predominantly from regions with well-established surveillance programs, thus potentially introducing geographic bias toward areas with greater resources or heightened concern about mosquito-borne diseases. Austria contributes 2,798 of the 5,129 weekly observations but only one non-zero record, so the climate–oviposition relationship is effectively learned from the Spanish, Albanian and Italian series; predictions for northern and eastern Europe therefore rest on few positive observations. Non-zero rates occur in 8 of the 13 countries and at 22 of the 75 monitoring sites, so stratified validation by country or landscape type is not supported by the present data. This may constrain model performance in under-sampled regions. For instance, the inability of our model to reproduce the secondary peak observed in a mountainous area of Italy may reflect an under-representation of such environments in the training dataset. *Second*, the reliance on egg counts as a proxy for adult abundance may introduce additional uncertainty. Egg-laying, egg-hatching rates and successful transition through the development stages to an adult may vary with local conditions and mosquito behaviour not fully explained by climate variables alone and may correlate with adult population sizes in complex ways. These limitations underscore the need for more comprehensive and harmonized surveillance efforts, ideally covering multiple years, seasons, diverse ecological contexts, and multiple mosquito life stages. The observedhistory Counter provides the highest predictive performance when recent trap measurements are available. For autonomous projections, however, such measurements are not available. Figure 7 therefore uses the dedicated autoregressive model, which was trained and evaluated using its own recursively generated previous predictions. These autonomous projections require further validation across additional regions, years, and surveillance systems before operational use. Models that can satisfactorily replicate observed mosquito population dynamics could then be combined with virus transmission models and disease importation data to inform public health about outbreak risks. Such integration is outside the scope of the present study, and the current egg-laying-rate outputs should not be interpreted as direct estimates of outbreak risk.

An important further limitation concerning the ***AI****edes* modelling approach (for both models) needs a mention. While neural networks deliver strong predictive performance, they provide limited interpretability, preventing straightforward identification of specific mechanistic phenomena, such as climate thresholds or factor interactions driving predictions. The input comparisons reported here quantify the contribution of variable groups through changes in held-out performance, but this is coarser than the explicit parameter estimates and effect sizes provided by traditional statistical techniques, and it does not resolve climate thresholds or interactions among predictors. Dedicated explainability analyses, such as SHAP values or permutation feature importance, together with hybrid data-mechanistic formulations, are a distinct line of work that we leave to future studies.

With this work, we demonstrate in practice the effectiveness of neural networks—an approach that is at present underutilised for the modelling of the geographical distribution of vectors, both for presence-absence classification and for abundance prediction. Our methodology is based on recent advances in the AI community [26], where benchmarking and standardized evaluation have accelerated progress across domains. To foster reproducibility and facilitate future comparisons, we release the code alongside ready-to-use datasets and detailed documentation, as a reference benchmark for the community [46, 33]. The logistic formulation used here is a functional-form reference rather than an exhaustive benchmark: it isolates the effect of adding hidden nonlinear layers to the same inputs, so our results do not establish that neural networks outperform other classes of distribution model. Systematic comparison with maximum-entropy models, generalised additive models, gradient-boosted trees and stacked ensembles, as well as with mechanistic life-cycle models, remains an open task that the released datasets, code, validation rules and reference metrics are intended to support. Our approach integrates multiple high-quality data sources to predict both the probability of presence and the weekly egg abundance of *Ae. albopictus* across Europe. With its diverse climatic conditions, different levels of mosquito establishment, and extensive surveillance programs, Europe provides a favourable setting for mosquito distribution modelling, yet, understandably, the methodology can be adapted to other regions, vector species, and to different spatial or temporal scales, provided adequate surveillance and environmental data are available.

Using two well-curated *Ae. albopictus* datasets, we have shown the promise of data-driven approaches based on neural networks in ecological modeling and prediction. We anticipate that, in the coming years, with the accumulation of new datasets, the adoption of more advanced neural network techniques, and the integration of additional input variables—such as satellite imagery—results could improve even further and more rapidly. Future work should focus on expanding the model validation to additional regions and surveillance systems, improving confidence in model generalizability. Incorporating higher-resolution spatial data and additional environmental variables could refine predictive accuracy, while the integration of explainable AI methods or hybrid data-mechanistic approaches could enhance interpretability. Ultimately, combining robust AI-driven predictive tools with sustained investment in entomological surveillance will be essential to strengthen vector monitoring and support effective public health preparedness and control of mosquito-borne infections.

## CRediT authorship contribution statement

Indaco Biazzo: Conceptualization, Methodology, Software, Formal analysis, Writing – original draft, Writing – review and editing.

Lia Orfei: Conceptualization, Methodology, Software, Formal analysis, Writing – original draft, Writing – review and editing.

Ioannis Kioutsioukis: Conceptualization, Formal analysis, Writing – review and editing. Lea Schuh: Conceptualization, Formal analysis, Writing – review and editing.

Céline Gossner: Formal analysis, Writing – review and editing. Olivier Briet: Formal analysis, Writing – review and editing.

Peter V. Markov: Supervision, Conceptualization, Formal analysis, Writing – original draft, Writing – review and editing.

## Declaration of Generative AI and AI-assisted technologies in the writing process

During the preparation of this work, the authors used GPT@JRC to support language polishing and improve the clarity of the manuscript. After using this tool, the authors reviewed and edited the content as needed and take full responsibility for the content of the published article.

## Data and code availability

The harmonised climate and *Aedes albopictus* surveillance datasets used in this study are publicly available through the European Commission, Joint Research Centre Data Catalogue [50], DOI: 10.2905/e4e68952-a02f-460d-8f85-eb630742741d. A detailed description of the dataset construction, harmonisation and validation is provided in the companion data-descriptor manuscript submitted to *Scientific Data* [33]. The source code, model implementation and documentation are available at [46]. The raw surveillance and climate data sources used to construct the datasets are cited in the Materials and methods section.

## Acknowledgements

We would like to thank the colleagues of the Digital Health Unit (JRC.F7) at the Joint Research Centre of the European Commission for their guidance and support.

The views expressed are those of the authors and may not in any circumstances be regarded as stating an official position of the European Commission.

## References

[1] R. C. Wilkerson, Y.-M. Linton, D. Strickman, Mosquitoes of the World, Johns Hopkins University Press, Baltimore, MD, USA, 2020.

[2] M. U. Kraemer, R. C. Reiner Jr, O. J. Brady, J. P. Messina, M. Gilbert, D. M. Pigott, D. Yi, K. Johnson, L. Earl, L. B. Marczak, et al., Past and future spread of the arbovirus vectors *Aedes aegypti* and *Aedes albopictus*, Nature Microbiology 4 (5) (2019) 854–863.

[3] S. J. Ryan, C. J. Carlson, E. A. Mordecai, L. R. Johnson, Global expansion and redistribution of aedes-borne virus transmission risk with climate change, PLoS Neglected Tropical Diseases 13 (3) (2019) e0007213. doi:10.1371/journal.pntd.0007213. URL https://journals.plos.org/plosntds/article?id=10.1371/journal.pntd.0007213

4. European Centre for Disease Prevention and Control and European Food Safety Authority, *Aedes albopictus* current known distribution: June 2025 (2025). URL https://ecdc.europa.eu/en/disease-vectors/surveillance-and-disease-data/mosquito-maps

[5] W. A. Hawley, The biology of *Aedes albopictus*, Journal of the American Mosquito Control Association 4 (2) (1988) 1–39.

[6] M. U. Kraemer, M. E. Sinka, K. A. Duda, A. Q. Mylne, F. M. Shearer, C. M. Barker, C. G. Moore, R. G. Carvalho, G. E. Coelho, W. Van Bortel, et al., The global distribution of the arbovirus vectors aedes aegypti and ae. albopictus, elife 4 (2015) e08347.

[7] G. Rezza, L. Nicoletti, R. Angelini, R. Romi, A. C. Finarelli, M. Panning, A. Cassone, Infection with chikungunya virus in italy: an outbreak in a temperate region, Lancet 370 (9602) (2007) 1840–1846.

[8] P. e. a. Cattaneo, Transmission of autochthonous aedes-borne arboviruses and related public health challenges in europe 2007–2023: a systematic review and secondary analysis, The Lancet Regional Health – Europe 51 (2023) 101231. doi:10.1016/j.lanepe.2025.101231.

[9] G. Guzzetta, F. Montarsi, F. A. Baldacchino, M. Metz, G. Capelli, A. Rizzoli, A. Pugliese, R. Rosà, P. Poletti, S. Merler, Potential risk of dengue and chikungunya outbreaks in northern italy based on a population model of aedes albopictus (diptera: Culicidae), PLoS Neglected Tropical Diseases 10 (6) (2016) e0004762. doi:10.1371/journal.pntd.0004762.

[10] I. Kioutsioukis, N. I. Stilianakis, Assessment of west nile virus transmission risk from a weather-dependent epidemiological model and a global sensitivity analysis framework, Acta Tropica 193 (2019) 129–141. doi:10.1016/j.actatropica.2019.03.003.

[11] R. Li, L. Xu, O. N. Bjørnstad, K. Liu, T. Song, A. Chen, B. Xu, Q. Liu, N. C. Stenseth, Climate-driven variation in mosquito density predicts the spatiotemporal dynamics of dengue, Proceedings of the National Academy of Sciences 116 (9) (2019) 3624–3629.

[12] A. Tran, M. Mangeas, M. Demarchi, E. Roux, P. Degenne, M. Haramboure, G. Le Goff, D. Damiens, L.-C. Gouagna, V. Herbreteau, et al., Complementarity of empirical and process-based approaches to modelling mosquito population dynamics with aedes albopictus as an example—application to the development of an operational mapping tool of vector populations, PloS one 15 (1) (2020) e0227407.

[13] A. Angelou, I. Kioutsioukis, N. I. Stilianakis, A climate-dependent spatial epidemiological model for the transmission risk of west nile virus at local scale, One Health 13 (2021) 100330. 10.1016/j.onehlt.2021.100330. URL https://www.sciencedirect.com/science/article/pii/S2352771421001208

[14] D. Da Re, W. Van Bortel, F. Reuss, et al., dynamaedes: a unified modelling framework for invasive *Aedes* mosquitoes, Parasites & Vectors 15 (2022) 414. doi:10.1186/s13071-022-05414-4. URL https://parasitesandvectors.biomedcentral.com/articles/10.1186/s13071-022-05414-4

[15] E. L. Davis, L. J. Reimer, L. Pellis, H. M. Lyons, An analytically tractable, agestructured model of the impact of vector control on mosquito-transmitted infections, PLoS Computational Biology 20 (1) (2024) e1011440.

[16] S. Barman, J. C. Semenza, P. Singh, H. Sjödin, J. Rocklöv, J. Wallin, A climate and population dependent diffusion model forecasts the spread of aedes albopictus mosquitoes in europe, Communications Earth & Environment 6 (1) (2025) 276. 10.1038/s43247-025-02199-z.

[17] N. Stilianakis, V. Syrris, T. Petroliagkis, P. Pärt, S. Gewehr, S. Kalaitzopoulou, S. Mourelatos, A. Baka, D. Pervanidou, J. Vontas, C. Hadjichristodoulou, Identification of climatic factors affecting the epidemiology of human west nile virus infections in northern greece, PLoS ONE 11 (09 2016). doi:10.1371/journal.pone.0161510.

[18] A. Zardini, F. Menegale, A. Gobbi, M. Manica, G. Guzzetta, V. d’Andrea, V. Marziano, F. Trentini, F. Montarsi, B. Caputo, et al., Estimating the potential risk of transmission of arboviruses in the americas and europe: a modelling study, The Lancet Planetary Health 8 (1) (2024) e30–e40.

[19] B. Yang, B. A. Borgert, B. W. Alto, C. K. Boohene, J. Brew, K. Deutsch, J. T. DeValerio, R. R. Dinglasan, D. Dixon, J. M. Faella, S. L. Fisher-Grainger, G. E. Glass, R. Hayes, D. F. Hoel, A. Horton, A. Janusauskaite, B. Kellner, M. U. G. Kraemer, K. J. Lucas, J. Medina, D. A. T. Cummings, Modelling distributions of aedes aegypti and aedes albopictus using climate, host density and interspecies competition, PLoS neglected tropical diseases 15 (3) (2021) e0009063. doi:10.1371/journal.pntd.0009063.

[20] A. Y. Lim, Y. Jafari, J. M. Caldwell, H. E. Clapham, K. A. M. Gaythorpe, L. HussainAlkhateeb, M. A. Johansson, M. U. G. Kraemer, R. J. Maude, C. P. McCormack, J. P. Messina, E. A. Mordecai, I. B. Rabe, R. C. Reiner, S. J. Ryan, H. Salje, J. C. Semenza, D. P. Rojas, O. J. Brady, A systematic review of the data, methods and environmental covariates used to map aedes-borne arbovirus transmission risk, BMC infectious diseases 23 (1) (2023) 708. doi:10.1186/s12879-023-08717-8.

[21] J. R. Barker, H. J. MacIsaac, Species distribution models applied to mosquitoes: Use, quality assessment, and recommendations for best practice, Ecological Modelling 472 (2022) 110073. doi:10.1016/j.ecolmodel.2022.110073. URL https://doi.org/10.1016/j.ecolmodel.2022.110073

[22] C. A. Lippi, S. J. Mundis, R. Sippy, et al., Trends in mosquito species distribution modeling: insights for vector surveillance and disease control, Parasites and Vectors 16 (2023) 302. doi:10.1186/s13071-023-05912-z.

[23] P. Georgiades, Y. Proestos, J. Lelieveld, K. Erguler, Machine learning modeling of aedes albopictus habitat suitability in the 21st century, Insects 14 (5) (2023) 447. doi: 10.3390/insects14050447.

[24] S. Taheri, M. A. González, M. J. Ruiz-López, R. Soriguer, J. Figuerola, Patterns of west nile virus vector co-occurrence and spatial overlap with human cases across europe, One Health 20 (2025) 101041. 10.1016/j.onehlt.2025.101041.

[25] D. Da Re, G. Marini, C. Bonannella, F. Laurini, M. Manica, N. Anicic, A. Albieri, P. Angelini, D. Arnoldi, F. Bertola, B. Caputo, C. De Liberato, A. Della Torre, E. Flacio, A. Franceschini, F. Gradoni, P. Kadriaj, V. Lencioni, I. Del Lesto, F. Russa, R. Rosà, Modelling the seasonal dynamics of *Aedes albopictus* populations using a spatio-temporal stacked machine learning model, Scientific Reports 15 (1) (2025) 3750. doi:10.1038/s41598-025-87554-y.

[26] M. U. G. Kraemer, J. L. H. Tsui, S. Y. Chang, et al., Artificial intelligence for modelling infectious disease epidemics, Nature 638 (2025) 623–635. doi:10.1038/s41586-024-08564-w. URL https://www.nature.com/articles/s41586-024-08564-w

[27] Q. Yuan, H. Shen, T. Li, Z. Li, S. Li, Y. Jiang, H. Xu, W. Tan, Q. Yang, J. Wang, J. Gao, L. Zhang, Deep learning in environmental remote sensing: Achievements and challenges, Remote Sensing of Environment 241 (2020) 111716. doi:10.1016/j.rse.2020.111716.

[28] E. A. Mordecai, J. M. Caldwell, M. K. Grossman, C. A. Lippi, L. R. Johnson, M. Neira, J. R. Rohr, S. J. Ryan, V. Savage, M. S. Shocket, R. Sippy, A. M. Stewart Ibarra, M. B. Thomas, O. Villena, Thermal biology of mosquito-borne disease, Ecology Letters 22 (10) (2019) 1690–1708. doi:10.1111/ele.13335.

[29] L. M. Beck-Johnson, W. A. Nelson, K. P. Paaijmans, A. F. Read, M. B. Thomas, O. N. Bjørnstad, The effect of temperature on *Anopheles* mosquito population dynamics and the potential for malaria transmission, PLoS One 8 (11) (2013) e79276. doi: 10.1371/journal.pone.0079276.

30. A. C. Kinney, R. Barrera, J. Lega, Rapid and accurate mosquito abundance forecasting with aedes-ai neural networks, arXiv preprint arXiv:2408.16152 (2024).

[31] M. Zhang, X. Wang, S. Tang, Integrating dynamic models and neural networks to discover the mechanism of meteorological factors on aedes population, PLOS Computational Biology 20 (9) (2024) e1012499.

[32] G. R. W. Wint, T. Balenghien, E. Berriatua, M. Braks, C. Marsboom, J. Medlock, F. Schaffner, W. Van Bortel, N. Alexander, B. Alten, E. Czwienczek, S. Dhollander, E. Ducheyne, C. M. Gossner, K. Hansford, G. Hendrickx, H. Honrubia, T. Matheussen, A. D. Mihalca, D. Petric, O. Briet, Vectornet: collaborative mapping of arthropod disease vectors in europe and surrounding areas since 2010, Euro surveillance: bulletin Européen sur les maladies transmissibles = European communicable disease bulletin 28 (26) (2023) 2200666. doi:10.2807/1560-7917.ES.2023.28.26.2200666.

33. I. Biazzo, S. Consoli, L. Orfei, L. Schuh, P. V. Markov, Harmonised climate and aedes albopictus arboviral vector mosquito surveillance datasets for the European continent, submitted to Scientific Data (2026).

34. ECDC, Aedes albopictus factsheet, available at: https://www.ecdc.europa.eu/en/disease-vectors/facts/mosquito-factsheets/aedes-albopictus (Accessed: 2025-03-11) (2024).

[35] M. Manica, R. Rosà, A. Della Torre, B. Caputo, From eggs to bites: do ovitrap data provide reliable estimates of aedes albopictus biting females?, PeerJ 5 (2017) e2998. doi:10.7717/peerj.2998.

[36] V. Steindorf, K. B., H. M., N. Stollenwerk, A. Cevidanes, J. F. Barandika, P. Vazquez, A. L. García-Pérez, M. Aguiar, Forecasting invasive mosquito abundance in the basque country, spain using machine learning techniques, Parasites & Vectors 18 (1) (2025) 109. doi:10.1186/s13071-025-06733-y.

[37] M. Á. Miranda, C. Barceló, D. Arnoldi, X. Augsten, K. Bakran-Lebl, G. Balatsos, M. Bengoa, P. Bindler, K. Boršová, M. Bourquia, D. Bravo-Barriga, V. Čabanová, B. Caputo, M. Christou, S. Delacour, R. Eritja, O. Fassi-Fihri, M. Ferraguti, E. Flacio, E. Frontera, et al., Aimsurv: First pan-european harmonized surveillance of aedes invasive mosquito species of relevance for human vector-borne diseases, GigaByte (Hong Kong, China) (2022). doi:10.46471/gigabyte.57. URL https://doi.org/10.46471/gigabyte.57

[38] VectorNet, European Centre for Disease Prevention and Control, European Food Safety Authority, Vectornet (2025). URL 10.15468/f3k8r9

[39] F. Romiti, R. Casini, A. Magliano, A. Ermenegildi, C. De Liberato, Aedes albopictus abundance and phenology along an altitudinal gradient in lazio region (central italy), Parasites & Vectors 15 (1) (2022) 92. 10.1186/s13071-022-05215-9.

40. European Commission, Copernicus: Europe’s eyes on Earth, the European Union’s Earth observation programme (2025). URL https://www.copernicus.eu/en

[41] D. P. van Vuuren, J. Edmonds, M. Kainuma, et al., The representative concentration pathways: an overview, Climatic Change 109 (2011) 5. doi:10.1007/s10584-011-0148-z.

42. WorldClim, Worldclim: Global climate data (2025). URL https://worldclim.org/data/index.html

[43] J. Muñoz Sabater, ERA5-Land hourly data from 1950 to present (2019). doi:10.24381/cds.e2161bac.

44. C. for Disease Control, Prevention, Aedes Mosquito Lifecycle (2025). URL https://www.cdc.gov/mosquitoes/pdfs/aedeslifecycle-p.pdf

45. A. Paszke, S. Gross, F. Massa, A. Lerer, J. Bradbury, G. Chanan, T. Killeen, Z. Lin, N. Gimelshein, L. Antiga, A. Desmaison, A. Kopf, E. Yang, Z. DeVito, M. Raison, A. Tejani, S. Chilamkurthy, B. Steiner, L. Fang, J. Bai, S. Chintala, Pytorch: An imperative style, high-performance deep learning library, in: Advances in Neural Information Processing Systems, Vol. 32, 2019.

46. Authors, Aiedes: Neural network models for predicting mosquito presence and abundance, https://github.com/AI4PHI/AIedes (2026).

[47] C. M. Bishop, Pattern Recognition and Machine Learning, Springer, 2006.

[48] D. P. Kingma, J. Ba, Adam: A method for stochastic optimization, arXiv preprint arXiv:1412.6980 (2014). URL https://arxiv.org/abs/1412.6980

[49] N. Srivastava, G. Hinton, A. Krizhevsky, I. Sutskever, R. Salakhutdinov, Dropout: A simple way to prevent neural networks from overfitting, Journal of Machine Learning Research 15 (56) (2014) 1929–1958.

50. I. Biazzo, L. Orfei, L. Schuh, S. Consoli, N. I. Stilianakis, P. V. Markov, Harmonised climate and aedes albopictus arboviral vector mosquito surveillance datasets for the european continent, dataset (2026). doi:10.2905/e4e68952-a02f-460d-8f85-eb630742741d. URL http://data.europa.eu/89h/e4e68952-a02f-460d-8f85-eb630742741d

