## Supplementary material for "AIedes: a neural-network framework for modelling the distribution and abundance of *Aedes albopictus* in Europe": SI

#### 1 Climate-Based Filtering of Distribution Data

As introduced in the Data and Methods section, under the subsection *Aedes albopictus datasets* and the paragraph *Mosquito presence prediction*, the ECDC/EFSA VectorNet maps [1, 2] provide the baseline presence and absence information for *Aedes albopictus* at the NUTS3 administrative level (NUTS2 for a few countries). To integrate these data with the Copernicus climate variables used in our model (spatial resolution of 12.5 km × 12.5 km), we reprojected the administrative-unit classifications onto the same regular grid as the Copernicus climate dataset, covering the European domain [3].

Such spatial downscaling may introduce artefacts, particularly spurious presences in climatically unsuitable areas, due to the coarse administrative boundaries relative to environmental gradients. To mitigate these effects, following an approach similar to [4], we applied a climate suitability mask to filter out implausible conditions.

Specifically, any grid cell classified as "present" after downscaling was reassigned to "absent" if local climate conditions fell outside the ecological thresholds reported in the ECDC guidelines [5], which define a more conservative climatic envelope than the one used by [4]. The mask combined two independent suitability functions:

- precipitation suitability, requiring annual precipitation greater than or equal to $200\,\mathrm{mm\,yr^{-1}}$;
- temperature suitability, requiring both a mean temperature of the coldest month greater than or equal to $-3$°C to ensure egg overwintering, and an annual mean temperature greater than or equal to 10°C to support development.

A grid cell was retained as suitable only if both precipitation and temperature criteria were met. This filtering step ensured internal consistency between the observed distribution and the climatic envelope of *Ae. albopictus*, thereby reducing the impact of administrative-level downscaling artefacts before model training.

#### 2 Geographic-transfer sensitivity analysis

##### 2.1 Motivation and NUTS3-level data splitting

The principal evaluation in this study measures predictive performance across the European domain represented by the available surveillance and climate data. A random division of grid cells addresses this objective because the model is intended to use information distributed across this observed domain. Nearby grid cells may share climatic and spatial characteristics, and this similarity is part of the interpolation setting addressed by the model.

As an additional sensitivity analysis, we examined geographic transfer by separating the data at the level of complete NUTS3 regions. All grid cells belonging to the same NUTS3 region were assigned to the same fold, and five-fold stratified grouped cross-validation was used to obtain predictions for every observation. Consequently, each region was evaluated using models trained without observations from that region.

This analysis addresses a different and more demanding prediction setting: transfer to geographic regions excluded from training. It is presented as a stress test of regional generalization rather than as a replacement for the principal evaluation. Because NUTS3 regions are administrative rather than climatic units, a withheld region may still share climatic conditions with regions represented during training. The analysis therefore evaluates geographic transfer rather than extrapolation to entirely unseen climatic conditions.

#### 2.2 Performance comparison across splitting strategies

Model performance was evaluated using accuracy, specificity, sensitivity, and critical success index (CSI). In each fold, predictions were evaluated only on observations from regions excluded from the corresponding training set. Five models with different random seeds were trained for each held-out fold, and their predictions were averaged separately for the no-hidden-layer logistic reference and the selected neural network. Across the five regional folds, the selected network had mean accuracy 0.920 $\pm$ 0.007, sensitivity 0.890 $\pm$ 0.048, specificity 0.936 $\pm$ 0.025, and CSI 0.797 $\pm$ 0.020. The corresponding logistic-reference values were 0.888 $\pm$ 0.020, 0.844 $\pm$ 0.035, 0.911 $\pm$ 0.038, and 0.728 $\pm$ 0.036. Performance was lower and more variable than in the random-split evaluation, as expected for this geographic-transfer setting. Nevertheless, the selected neural network maintained its improvement over the logistic reference for all four metrics.

#### 2.3 Spatial predictions under NUTS3-level splitting

Figure 1 reproduces the spatial comparison shown in the main text, but using predictions from models trained under the NUTS3-level split. Panel (a) shows predictions from the simplest neural network architecture, equivalent to logistic regression, while panel (b) shows predictions from the selected deep neural network, featuring three hidden layers with 50 units each. Panel (c) summarizes performance using predictions for observations from regions excluded from the corresponding training fold.

The regional separation produces more false-positive and false-negative classifications than the random split, particularly for the selected neural network, while the neural network continues to outperform the logistic reference under the same held-region evaluation.

#### 2.4 Summary

This sensitivity analysis shows that the random split and the NUTS3 split answer different questions. The random split measures interpolation within the geographic and climatic domain represented in the data, whereas the NUTS3 split measures transfer to administrative regions excluded from training. The lower and more variable NUTS3 metrics indicate that geographic transfer is more challenging than in-domain interpolation. At the same time, the improvement of the selected neural network over the logistic reference is preserved under regional separation.

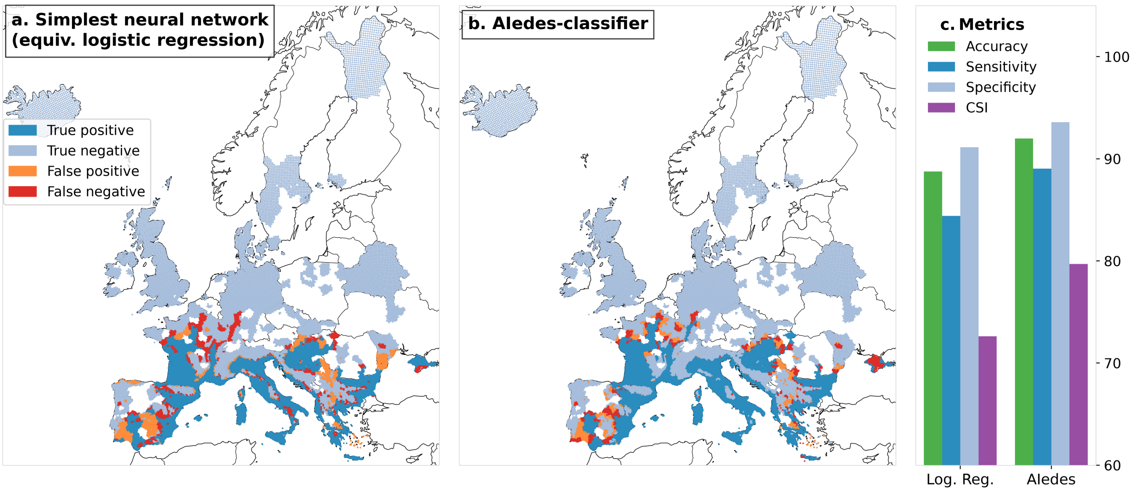


Figure 1: **Classification outcomes for *Ae. albopictus* under NUTS3-level train–test separation.** **(a)** Held-region results from the simplest neural network architecture tested, featuring no hidden layers and statistically equivalent to logistic regression. **(b)** Held-region results from the selected ***AI****edes*-Classifier, consisting of three hidden layers with 50 units each. **(c)** Pooled accuracy, sensitivity, specificity and critical success index (CSI) across all held-region predictions.

#### 3 Comparison with WorldClim-Based Distribution Modelling

In the main text, we compared the performance of the `AIedes`-classifier with that of a logistic regression model, i.e. the same functional form adopted by [4], but applied to different climatic input datasets.

The model by [4] was developed using the *WorldClim* bioclimatic dataset [6], which provides long-term averaged climatic conditions for the period 1970–2000. In contrast, our analyses rely on Copernicus-based climate variables, which better reflect the current environmental conditions corresponding to the mosquito records collected in 2023. Moreover, Copernicus products offer greater flexibility for future predictive applications, given their wide range of climate projection models and continuous availability for upcoming years.

In [4], four WorldClim variables were used as key climatic predictors: annual mean temperature, maximum temperature of the warmest month, annual precipitation, and precipitation of the warmest quarter (available at `https://www.worldclim.org/`). To ensure a consistent comparison, we repeated our analysis using these same WorldClim variables.

Figure 2 compares our predicted distribution of *Aedes albopictus* presence with the reference logistic-regression predictions based on the formulation used by [4]. The resulting classification maps and performance metrics show that the improvement achieved by our framework is preserved when it is applied using the original WorldClim-based input variables.

#### 4 Evaluation of Abundance Predictions

In the Results section of the main text, we reported the performance of the `AIedes`-counter in predicting mosquito abundance, specifically the weekly egg-laying rates. In the corresponding scatter plots shown in the main text, the model appears to slightly underpredict the observed values, particularly at higher egg counts. To determine whether this apparent deviation reflects a systematic bias or falls within the expected range of natural variability, we performed a Bland–Altman analysis on the held-out-trap predictions (Fig. 3).

The Bland–Altman plot compares the differences between predicted and observed egg counts

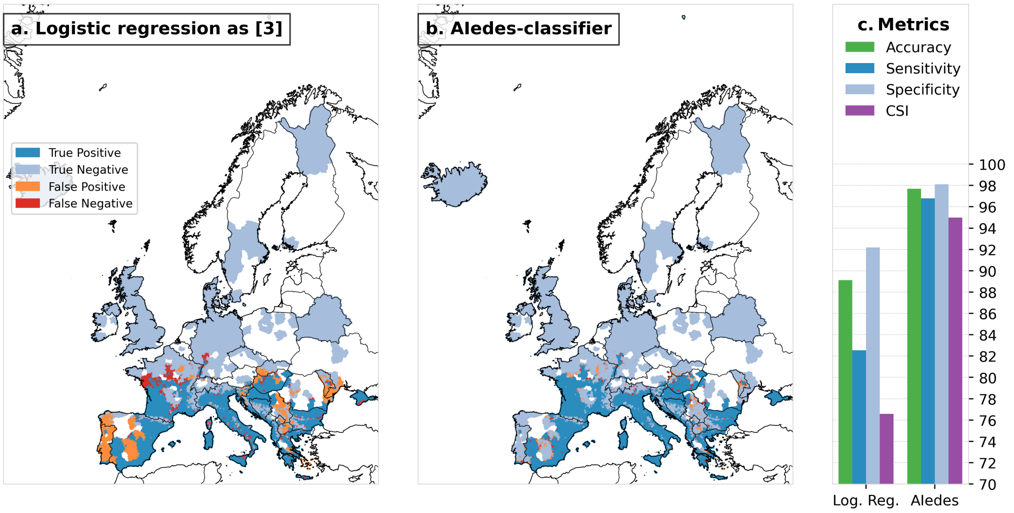


Figure 2: Comparison between the predicted presence maps produced by the `AIedes`-classifier (right) and the reference logistic-regression predictions based on the formulation used by [4] (left). The classification maps and performance metrics show that the improvement identified in the main analysis is preserved when the original WorldClim-based climatic variables are employed.

against their mean values, thus enabling a direct evaluation of agreement beyond simple correlation metrics. The mean difference between predictions and observations was close to zero ($-2.11$), indicating little overall mean bias. The 95% limits of agreement ($\pm 1.96$ SD) ranged from approximately $-53$ to $+49$, capturing the natural variability inherent in oviposition data.

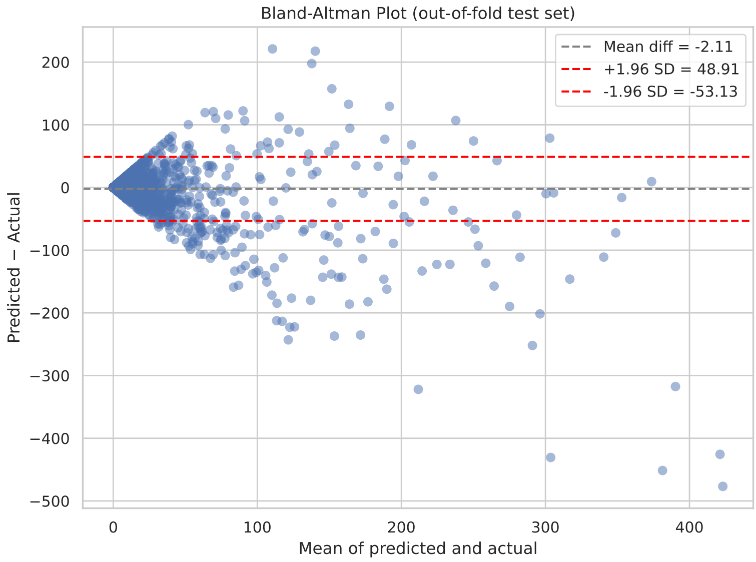


Figure 3: Bland–Altman plot comparing predicted and observed egg counts for traps excluded from the corresponding training sets. The mean difference ($-2.11$) indicates little overall mean bias, while the 95% limits of agreement (dashed lines) reflect variability consistent with the stochasticity of oviposition data.

### 5 Counter Ablation: Inputs, Architectures, Dropout, Loss Functions and Climate-History Windows

This section reports the complete sensitivity screen of the `AIedes`-counter. Every configuration that completed training is reported; no run was discarded on the basis of its performance, and no top-fraction filtering was applied.

#### 5.1 Screening design

Configurations were evaluated with the five-fold grouped cross-validation by trap described in the main text, so every evaluated observation comes from a trap excluded from the corresponding training set. The screen combined

- three loss functions: *Weighted Mean Squared Error* (WMSE), *Weighted Mean Squared Logarithmic Error* (WMSLE), and a zero-inflated-negative-binomial-inspired objective (ZINB);
- three dropout rates: 0.0, 0.2, 0.4;
- four hidden-layer architectures: [32], [64], [32, 16], and [64, 32];
- the input sets described below.

Each configuration was trained once in each of the five folds, giving 516 configurations and 2,580 retained training runs. Training used a batch size of 64, at most 3,000 epochs and a patience of 200 epochs, with loss-specific learning rates chosen for training stability: $1 \times 10^{-4}$ for WMSE, $1 \times 10^{-3}$ for WMSLE, and $5 \times 10^{-4}$ for ZINB. All reported values are means $\pm$ standard errors across the five trap-disjoint folds.

The input sets use the variable notation defined in the Data section of the main text. Two climate families were evaluated, $(T, P)$ and $(T, P, D)$, each represented either as daily values or as weekly averages over the same history. Each climate family was evaluated with and without the previous egg-laying rates, abbreviated $(E, I)$, which denote $E_{1w}$, $E_{2w}$ and their availability indicators $I_{1w}$, $I_{2w}$. This gives the eight climate-based input sets shown in Figures 4 and 5 for the 90-day history, represented either as 90 daily values ($90d$) or as 13 weekly averages ($13w$). Two further groups complete the screen: configurations that use $(E, I)$ alone, without any climate input, which occupy the first position of the same two figures, and, for WMSLE, the shorter 30- and 60-day histories (Figure 6).

#### 5.2 Loss functions, dropout and architectures

Figures 4 and 5 report held-out-trap $R^2$ and zero/non-zero accuracy for all nine input sets; the eight climate-based sets are discussed here and the no-climate set in Section 5.3. Three patterns are consistent across the three losses among the climate-based sets. First, adding the previous egg-laying rates raises the mean $R^2$ of the corresponding climate-only input set, from 0.460 to 0.507 for WMSE, from 0.441 to 0.498 for WMSLE, and from 0.463 to 0.499 for ZINB. Second, the effect of the temporal representation depends on the loss: WMSLE favours the daily representation (0.497 versus 0.442 on average), whereas WMSE slightly favours weekly averages (0.498 versus 0.469) and ZINB is insensitive to it (0.481 in both cases). Third, the differences among the four architectures are small relative to the fold-to-fold standard errors, and they do not identify a single best network.

Dropout acts differently on the two metrics. A rate of 0.2 gives the highest mean $R^2$ for all three losses (0.491 for WMSE, 0.481 for WMSLE, 0.486 for ZINB), while zero/non-zero accuracy decreases monotonically with dropout, from 0.895 to 0.871 for WMSE, from 0.921 to 0.916 for WMSLE, and from 0.916 to 0.909 for ZINB. The accuracy differences are small for WMSLE and ZINB and considerably larger for WMSE.

Comparing losses, WMSLE and ZINB classify zeros and non-zeros more reliably than WMSE. Across the 96 climate-based configurations of each loss, accuracy never falls below 0.888 for WMSLE and 0.871 for ZINB, with only one and ten configurations below 0.90 respectively, whereas WMSE falls to 0.807 and lies below 0.90 in 63 of its 96 configurations.

The three losses reach comparable $R^2$ values on the original scale, with overlapping standard errors, so this metric alone does not discriminate between them. They are clearly separated, however, on the logarithmic scale, which is the relevant scale for a distribution in which about 80% of the weekly rates are zero and the non-zero values are strongly right-skewed. Averaged over the 96 climate-based configurations of each loss, log-$R^2$ is 0.661 for WMSLE, 0.602 for ZINB and 0.503 for WMSE, and the weakest WMSLE configuration (0.619) still exceeds the mean of both alternatives. WMSLE was therefore selected for the main analysis: it is the only loss that is simultaneously best on the logarithmic scale and best at separating zero from non-zero weeks, while remaining statistically indistinguishable from the alternatives on the original scale.

#### 5.3 Configurations using only previous egg-laying rates

The first position of Figures 4 and 5, labelled $(E, I)$, reports the configurations that receive the previous egg-laying rates alone. Their best mean $R^2$ is $0.432 \pm 0.032$ for WMSE, $0.359 \pm 0.020$ for WMSLE and $0.436 \pm 0.032$ for ZINB, all below the corresponding climate-based configurations. Their zero/non-zero accuracy is both lower and far more variable, ranging from 0.20 to 0.83: several of these models never predict a zero and therefore reproduce only the non-zero fraction of the data. Because the vertical range of both figures is set by the climate-based configurations, all 36 of these configurations are visible in Figure 4, whereas 33 of them lie below the accuracy range of Figure 5: five appear there as truncated intervals at the lower edge and the remaining 28 are outside the panel. Previous egg-laying rates alone are thus insufficient, and they are informative only in combination with the climate history.

#### 5.4 Length of the climate history

Figure 6 compares 30-, 60- and 90-day climate histories for the WMSLE loss, which is the loss used in the main analysis; the 30- and 60-day histories were run for this loss only. Longer histories improve performance for every input set, and the gain from 60 to 90 days is smaller than the gain from 30 to 60 days. The effect is strongest for weekly averages of $(T, P)$ without previous rates, which rise from $0.208 \pm 0.020$ at 30 days to $0.386 \pm 0.011$ at 90 days, and weakest for daily $(T, P, D)$ with previous rates, which rise from $0.498 \pm 0.023$ to $0.523 \pm 0.023$. Daily representations dominate weekly averages at every window length, and the two converge as the window grows. Windows longer than 90 days were not evaluated because the published surveillance dataset provides exactly 90 days of climate history for each observation.

#### 5.5 Number of previous weekly rates

The screen above always supplies the previous rates in pairs, $E_{1w}$ and $E_{2w}$. To isolate the contribution of the second week, the selected WMSLE configuration—daily $(T, P, D)_{90d}$, a [64] network and dropout 0.2—was retrained with one previous week only, keeping every other setting and the same folds and seeds. With a single previous week the mean held-out-trap $R^2$ is $0.529 \pm 0.033$, against $0.553 \pm 0.023$ with two previous weeks; the corresponding log-$R^2$ values are $0.673 \pm 0.016$ and $0.670 \pm 0.013$, RMSE $26.8 \pm 0.9$ and $26.2 \pm 0.9$, and zero/non-zero accuracy $0.922 \pm 0.004$ and $0.923 \pm 0.004$. Comparing the two lag orders fold by fold, the paired difference (one week minus two weeks) is $-0.024 \pm 0.018$ in $R^2$, $+0.003 \pm 0.005$ in log-$R^2$, $+0.62 \pm 0.50$ in RMSE and $-0.0015 \pm 0.0010$ in accuracy. The second previous week therefore gives a small average gain on the original scale that is not resolved by the fold-to-fold variability, and no loss

Table 1: Best-performing configuration per loss function, selected by mean held-out-trap $R^2$ in the screen and then retrained with five independent seeds in each of the five trap-disjoint folds. Entries report means $\pm$ standard errors across folds after averaging seeds within each fold. $(E, I)$ denotes the previous egg-laying rates $E_{1w}$ and $E_{2w}$ with their availability indicators $I_{1w}$ and $I_{2w}$; DO/Arch gives the dropout rate and the hidden-layer widths. Log-$R^2$ uses the $\log(1 + x)$ transformation and is the criterion, together with the zero/non-zero accuracy, on which WMSLE was preferred.

| LossFn | $R^2$ | Log-$R^2$ | Acc | RMSE | DO/Arch | Inputs |
|---|---|---|---|---|---|---|
| WMSE | $0.564 \pm 0.034$ | $0.503 \pm 0.019$ | $0.884 \pm 0.002$ | $25.8 \pm 0.9$ | 0.2/[64] | $(T, P)_{13w} + (E, I)$ |
| WMSLE | $0.553 \pm 0.023$ | $0.670 \pm 0.013$ | $0.923 \pm 0.004$ | $26.2 \pm 0.9$ | 0.2/[64] | $(T, P, D)_{90d} + (E, I)$ |
| ZINB | $0.527 \pm 0.023$ | $0.611 \pm 0.013$ | $0.905 \pm 0.003$ | $27.0 \pm 1.2$ | 0.2/[32, 16] | $(T, P)_{13w} + (E, I)$ |

on the logarithmic scale. We retain two previous weeks in the main analysis, which also covers the two weeks preceding the observation, consistent with the egg-to-adult development time.

##### 5.6 Best configuration per loss function

Table 1 reports, for each loss, the configuration with the highest mean held-out-trap $R^2$ in the screen, retrained with five independent seeds in each fold. On the original scale the three losses differ by about one standard error in $R^2$, and WMSE attains the lowest RMSE, so the raw-scale metrics give no decisive argument. The ordering is unambiguous on the logarithmic scale, where WMSLE reaches $0.670 \pm 0.013$ against $0.611 \pm 0.013$ for ZINB and $0.503 \pm 0.019$ for WMSE, and in zero/non-zero accuracy, where WMSLE reaches $0.923 \pm 0.004$ against $0.905 \pm 0.003$ and $0.884 \pm 0.002$. These two criteria motivate the WMSLE configuration used in the main text.

#### 6 Architecture Exploration for the Dedicated Autoregressive Counter

The configurations discussed above receive the measured previous egg-laying rates, so their performance describes prediction with continuing trap observations. For projections that proceed without further observations, the Counter must supply its own previous values, and the main text reports this autonomous mode separately in its comparison of Counter prediction modes. The dedicated autoregressive model is trained directly in that regime: during training the inputs $E_{1w}$ and $E_{2w}$ are the model's own predictions from one and two weeks earlier, generated recursively from a zero-initialised sequence, and the climate inputs remain the daily histories $T_{90d}$, $P_{90d}$ and $D_{90d}$ with the WMSLE loss and dropout 0.2.

Because recursive training is considerably more expensive than teacher-forced training, the architecture choice was made with a focused exploration rather than with the full grid. Two networks were compared, [64] and [64, 32], each trained once per fold on the same five trap-disjoint folds with the recursive rollout. At the last recorded checkpoint of these exploration runs, which lies between epochs 520 and 560 in every fold, the rollout $R^2$ on the held-out traps is $0.469 \pm 0.014$ for [64] and $0.493 \pm 0.031$ for [64, 32].

Both architectures were then trained to the full budget of 3,000 epochs with five independent initialisations in each fold, giving 25 models per architecture. Table 2 reports the resulting out-of-fold seed-ensemble predictions on the recursive cohort of 5,036 observations from 375 traps, the same cohort used for the prediction-mode comparison in the main text. The [64, 32] network gives the higher original-scale $R^2$ and the lower RMSE, with log-$R^2$ and zero/non-zero accuracy practically unchanged, and it is therefore the configuration used for the autonomous projections in the main text.

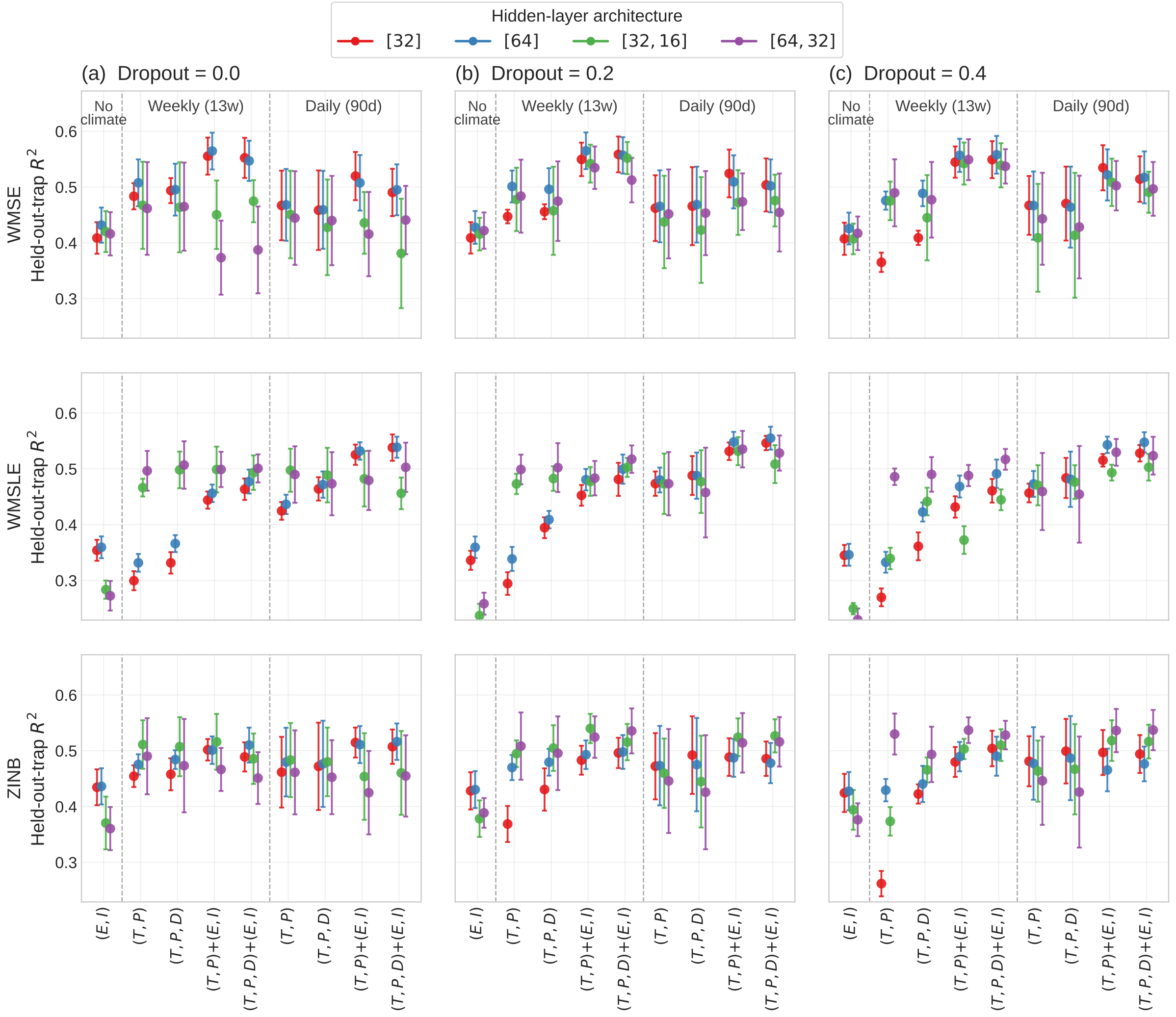


Figure 4: Held-out-trap $R^2$ across input sets, dropout rates and loss functions. Rows correspond to the loss functions and columns to the dropout rates. Within each panel the first position holds the configurations without climate inputs, $(E, I)$, and the remaining eight are the climate-based input sets for the 90-day history, represented as 13 weekly averages ($13w$) or 90 daily values ($90d$); dashed lines separate the three groups. $(E, I)$ denotes $E_{1w}$, $E_{2w}$, $I_{1w}$ and $I_{2w}$. Colours identify the four hidden-layer architectures, points are means across the five trap-disjoint folds and error bars are the corresponding standard errors. The vertical range is set by the climate-based configurations. Each configuration was trained once per fold and every completed run is retained.

Table 2: Dedicated autoregressive Counter: comparison of the two explored hidden-layer architectures. Both use the daily climate histories $T_{90d}$, $P_{90d}$ and $D_{90d}$, recursively generated $E_{1w}$ and $E_{2w}$, the WMSLE loss and dropout 0.2. Each entry is computed from the out-of-fold seed-ensemble predictions of 25 models (five folds × five initialisations) on the common recursive cohort of 5,036 observations from 375 traps. Log-$R^2$ uses the $\log(1+x)$ transformation; accuracy is the zero versus non-zero agreement.

| Architecture | $R^2$ | Log-$R^2$ | RMSE | Accuracy |
|---|---|---|---|---|
| [64] | 0.521 | 0.660 | 27.68 | 0.920 |
| [64, 32] | 0.535 | 0.663 | 27.25 | 0.919 |

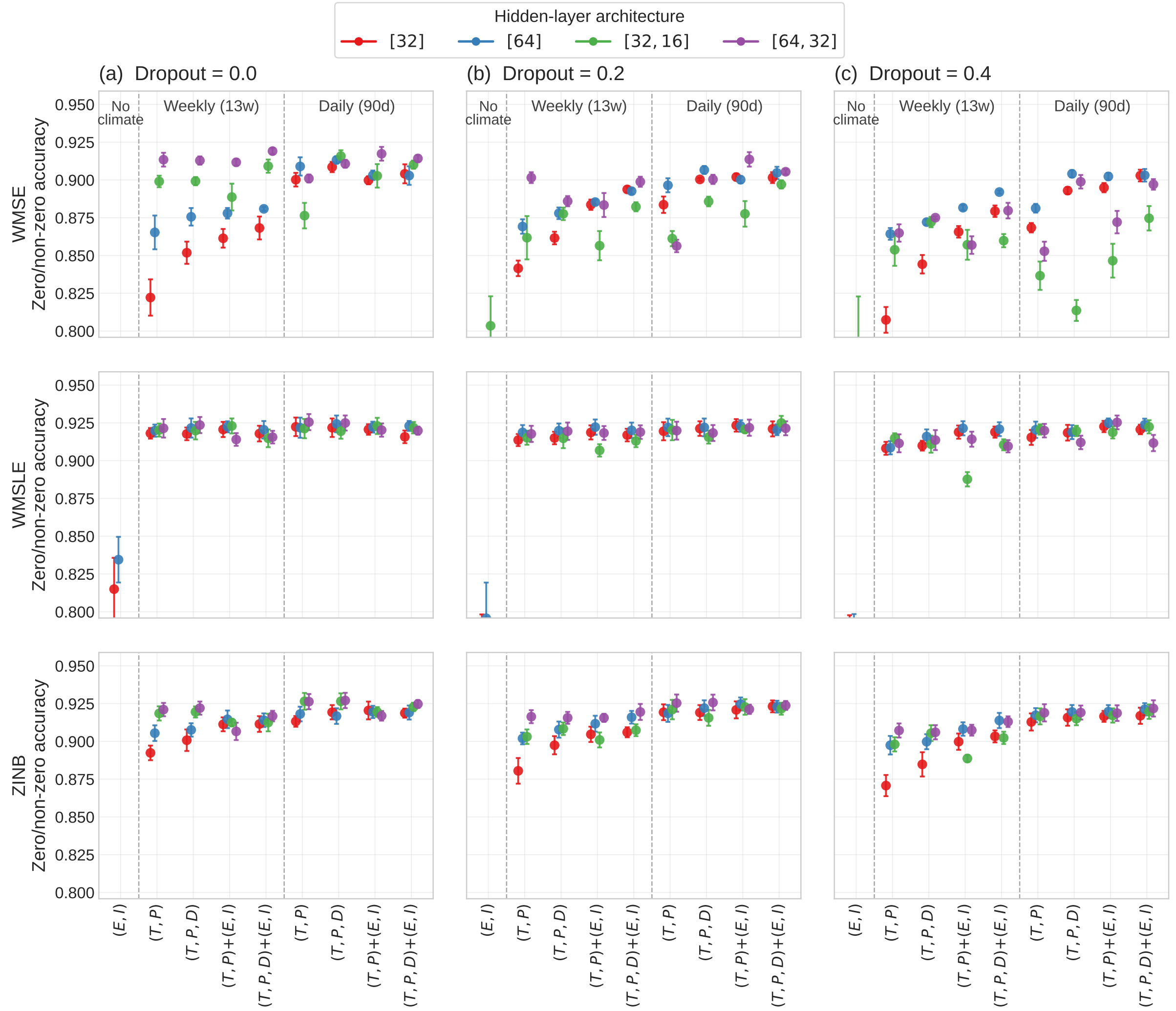


Figure 5: Zero/non-zero accuracy for the same configurations as Figure 4. WMSLE and ZINB classify zero and non-zero weekly rates more reliably than WMSE across all input sets, dropout rates and architectures. As in Figure 4, the vertical range is set by the climate-based configurations; several $(E, I)$ configurations reach accuracies as low as 0.20, because they never predict a zero weekly rate, and therefore lie below the plotted range. Points are means across the five trap-disjoint folds and error bars are the corresponding standard errors.

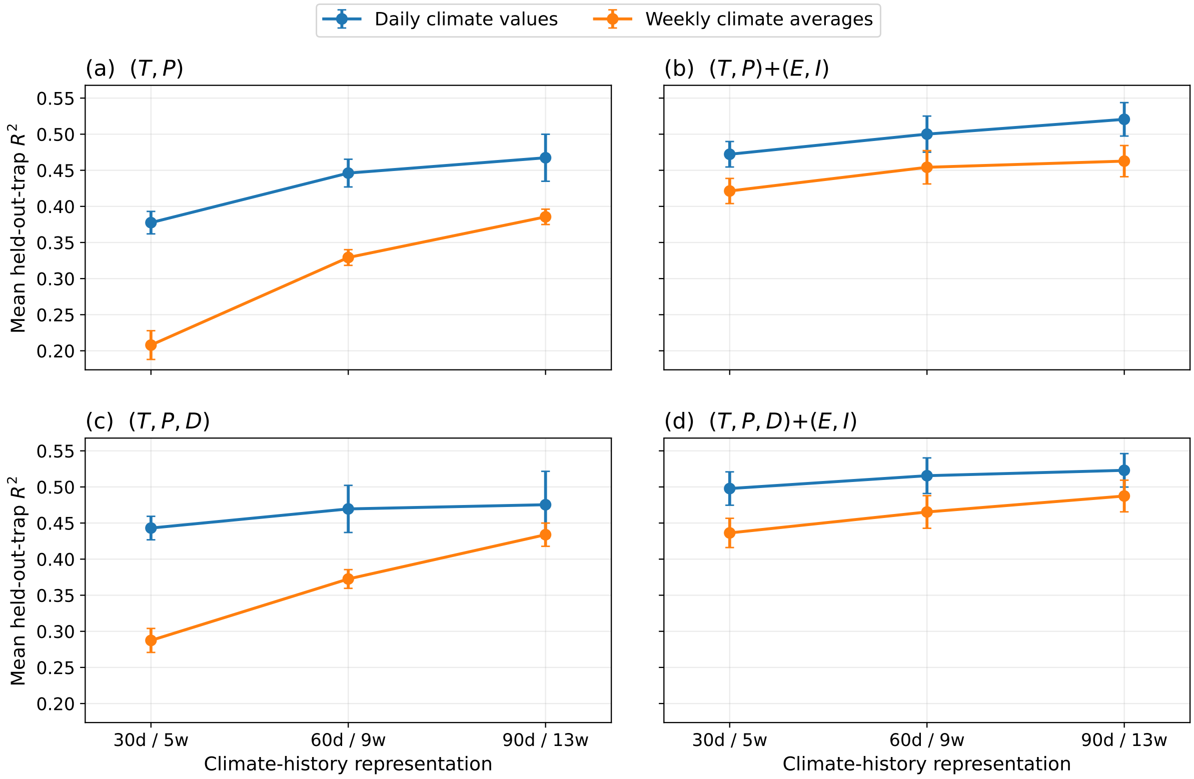


Figure 6: Sensitivity to the length of the climate history for the WMSLE loss. Each panel corresponds to one input set, with $(E, I)$ denoting $E_{1w}$, $E_{2w}$, $I_{1w}$ and $I_{2w}$, and compares daily climate values with weekly averages over the same 30-, 60- and 90-day windows; the tick labels give both representations of each window. Within each fold, the four architectures and three dropout rates are averaged with equal weight; points are means across the five trap-disjoint folds and error bars are the corresponding standard errors.

#### References


[1] European Centre for Disease Prevention and Control and European Food Safety Authority. *Aedes albopictus - current known distribution: June 2025*. Stockholm, 2025. URL: https://ecdc.europa.eu/en/disease-vectors/surveillance-and-disease-data/mosquito-maps.

[2] G. R. W. Wint et al. "VectorNet: collaborative mapping of arthropod disease vectors in Europe and surrounding areas since 2010". In: *Euro surveillance: bulletin Européen sur les maladies transmissibles = European communicable disease bulletin* 28.26 (2023), p. 2200666. DOI: 10.2807/1560-7917.ES.2023.28.26.2200666.

[3] Indaco Biazzo et al. "Harmonised climate and *Aedes albopictus* arboviral vector mosquito surveillance datasets for the European continent". Submitted to *Scientific Data*. 2026.

[4] Agnese Zardini et al. "Estimating the potential risk of transmission of arboviruses in the Americas and Europe: a modelling study". In: *The Lancet Planetary Health* 8.1 (2024), e30–e40.

[5] ECDC. *Aedes albopictus Factsheet*. Available at: https://www.ecdc.europa.eu/en/disease-vectors/facts/mosquito-factsheets/aedes-albopictus (Accessed: 2025-03-11). 2024.

[6] WorldClim. *WorldClim: Global Climate Data*. 2025. URL: https://worldclim.org/data/index.html (visited on 05/05/2025).
